# A Systematic Evaluation of Single-cell RNA-sequencing Imputation Methods

**DOI:** 10.1101/2020.01.29.925974

**Authors:** Wenpin Hou, Zhicheng Ji, Hongkai Ji, Stephanie C. Hicks

## Abstract

The rapid development of single-cell RNA-sequencing (scRNA-seq) technology, with increased sparsity compared to bulk RNA-sequencing (RNA-seq), has led to the emergence of many methods for preprocessing, including imputation methods. Here, we systematically evaluate the performance of 18 state-of-the-art scRNA-seq imputation methods using cell line and tissue data measured across experimental protocols. Specifically, we assess the similarity of imputed cell profiles to bulk samples as well as investigate whether methods recover relevant biological signals or introduce spurious noise in three downstream analyses: differential expression, unsupervised clustering, and inferring pseudotemporal trajectories. Broadly, we found significant variability in the performance of the methods across evaluation settings. While most scRNA-seq imputation methods recover biological expression observed in bulk RNA-seq data, the majority of the methods do not improve performance in downstream analyses compared to no imputation, in particular for clustering and trajectory analysis, and thus should be used with caution. Furthermore, we find that the performance of scRNA-seq imputation methods depends on many factors including the experimental protocol, the sparsity of the data, the number of cells in the dataset, and the magnitude of the effect sizes. We summarize our results and provide a key set of recommendations for users and investigators to navigate the current space of scRNA-seq imputation methods.

## 1 Introduction

Recent advances in high-throughput technologies have been developed to measure gene expression in individual cells^1–5^. In contrast to bulk RNA-sequencing (RNA-seq), a distinctive feature of data measured using single-cell RNA-sequencing (scRNA-seq) is the increased sparsity, or fraction of observed ‘zeros’, where a zero refers to no unique molecular identifiers (UMIs) or reads mapping to a given gene in a cell^6–9^. These observed zeros can be due to biological (relevant or nuisance) fluctuations in the measured trait or technical limitations related to challenges in quantifying small numbers of molecules. Examples of the latter include mRNA degradation during cell lysis or variation by chance of sampling lowly expressed transcripts^10^. The word *dropout*^6–8^ has been previously used to describe both biological and technical observed zeros, but the problem with using this catch-all term is it does not distinguish between the types of sparsity^10^.

To address the increased sparsity observed in scRNA-seq data, recent work has led to the development of “imputation” methods, in a similar spirit to imputing genotype data for genotypes that are missing or not observed. However, one major difference is that in scRNA-seq standard transcriptome reference maps, such as the Human Cell Atlas^11^ or the Tabula Muris Consortium^12^ are not yet widely available for all species, tissue types, genders, and so on. Therefore, the majority of imputation methods developed to date do not rely on an external reference map. They can also be categorized into three broad approaches^10^. The first group are imputation methods that directly model the sparsity using probabilistic models. These methods may or may not distinguish between biological and technical zeros, but if they do, they typically impute gene expression values for only the latter. A second approach adjusts (usually) all values (zero and non-zero) by smoothing or diffusing the raw expression values of cells with a similar expression profiles identified, for example, using neighbors in graph. The third approach first identifies a latent space representation of the cells, either through low-rank matrix-based methods (capturing linear relationships) or deep-learning methods (capturing non-linear relationships), and then reconstructs the observed expression matrix from the low-rank or estimated latent spaces, which will no longer be sparse. For the deep-learning approaches, such as variational autoencoders, both the estimated latent spaces and the “imputed” data (i.e. reconstructed expression matrix) can be used for downstream analyses, but otherwise only the imputed data is typically provided for downstream analyses.

Due to their recent and concurrent development, evaluations and comparisons between scRNA-seq imputation methods have been limited or restricted to a subset of imputation methods and downstream applications^13–16^. Furthermore, imputation methods can require varying types of raw or processed data as input, may rely on different methodological assumptions, and may be appropriate for only certain scRNA-seq experimental protocols, such UMI-based^1–3^ or full-length^4, 17^ transcript methods. Given these differences, the performance of these methods has been shown to vary in the evaluations to-date. For example, one study found imputation methods can introduce false signals when identifying differentially expressed genes with model-based methods outperforming smoothing-based methods, in particular for genes with a small effect size (log2 fold-change)^14^. Another study found imputation methods can introduce spurious correlations between imputed expression and total UMI counts^15^. Alternatively, others have shown spurious structural patterns in low dimensional representations of imputed data^14, 18^, which we also find in data where we expect no structural patterns in the data but patterns associated with library size emerge in the imputed data (Figures 1A, S1). In contrast, others have found a subset of imputation methods to be helpful to estimate library size factors for normalization of sparse scRNA-seq data^16^. Therefore, the answer to the question of which methods *can*, let alone *should*, be used for a particular analysis is often unclear.

**Figure 1.**
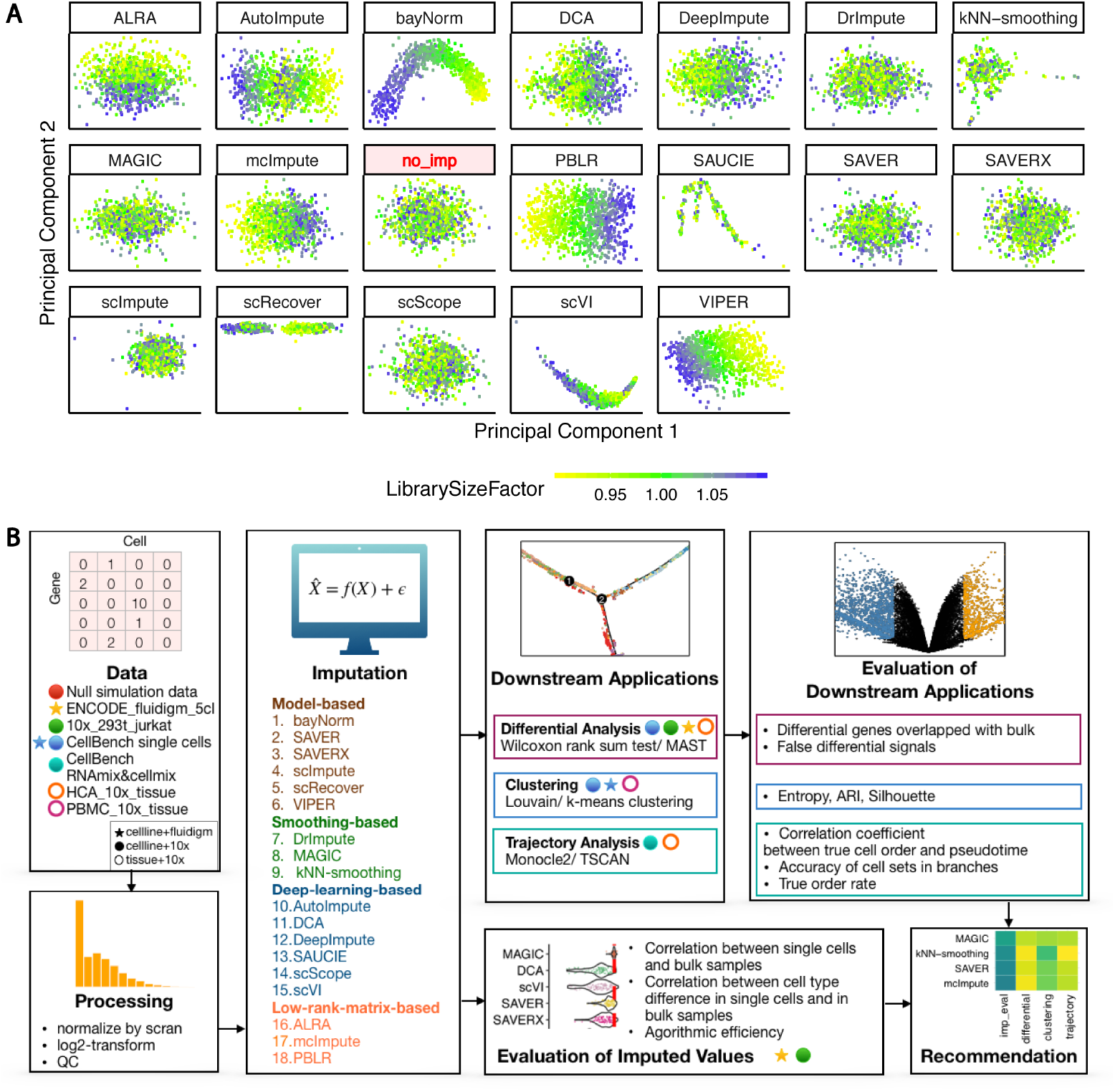
Motivation and overview of benchmark evaluation of scRNA-seq imputation methods. **(A)** Dimension reduction results after applying Principal Components Analysis (PCA) from either no imputation method (*no_imp* highlighted in red) or the 18 imputation methods using the *null simulations* data (Section 5.3) in which no structural pattern is expected. The color represents the simulated library size (defined as the total sum of counts across all relevant features) for each cell. **(B)** An overview of the benchmark comparison evaluating 18 scRNA-seq imputation methods.

To address this gap, we performed a systematic benchmark comparison and evaluation of 18 state-of-the-art scRNA-seq imputation methods (Figure 1B). Specifically, we evaluated (1) model-based imputation methods [bayNorm^19^, SAVER^20^, SAVER-X^21^, scImpute^22^, scRecover^23^, VIPER^24^], (2) smooth-based imputation methods [DrImpute^25^, MAGIC^26^, kNN-smoothing^27^], and (3) data reconstruction methods either using deep-learning methods [AutoImpute^28^, DCA^29^, DeepImpute^30^, SAUCIE^31^, scScope^32^, scVI^33^] or low-rank matrix-based methods [ALRA^34^, mcImpute^35^, PBLR^36^]. Throughout, we use `teletype` (or `monospace`) font when referring to specific software packages and *italicized* font when referring to datasets. While many of the imputation methods used raw scRNA-seq UMI or read counts as input, a subset of these methods required normalized (or log-transformed normalized) counts. In the latter case, we used the scRNA-seq pooling normalization method^37^ implemented in the `scran`^38^ R/Bioconductor^39, 40^ package, which has been previously shown to outperform other scRNA-seq normalization methods in full-length and UMI based methods^16, 18^. We also included a baseline comparison of “no imputation”, which is the raw scRNA-seq counts that have been adjusted for library size using the `scran`^37^ normalization method. While there are more methods available that could be used for imputation, we only included methods in our benchmark that (i) were originally designed (or specified by the authors) to be used as an imputation method, (ii) included software for users to download and run locally, and (iii) did not need external pieces of information (e.g. a network or an external reference map) and used scRNA-seq counts (raw or normalized) as input.

In this paper, we first evaluate the performance of the imputation method themselves on their ability to recover true expression values by comparing the similarity between imputed cell profiles and bulk samples in a homogeneous population of cells. Then, we investigate the performance of the imputation methods in downstream analyses including differential expression analysis, unsupervised clustering, and trajectory analysis. In addition to simulated data, we used two types of real single-cell data: cell lines and tissues measured across experimental platforms, including full-length and UMI methods using plate-based and droplet-based protocols. We summarize our results and provide a key set of recommendations for users and investigators to navigate the current space of scRNA-seq imputation methods.

## 2 Results

### 2.1 Similarity between bulk RNA-seq and imputed scRNA-seq data

To evaluate an imputation method’s ability to recover biological expression observed in bulk RNA-seq data, we assessed the similarity between bulk and imputed scRNA-seq data using cell lines (Figure 2). For this initial test, we focused on cell lines since they are less heterogeneous than tissues and have well-defined bulk expression profiles. The imputed values from the scRNA-seq data were evaluated using two approaches.

**Figure 2.**
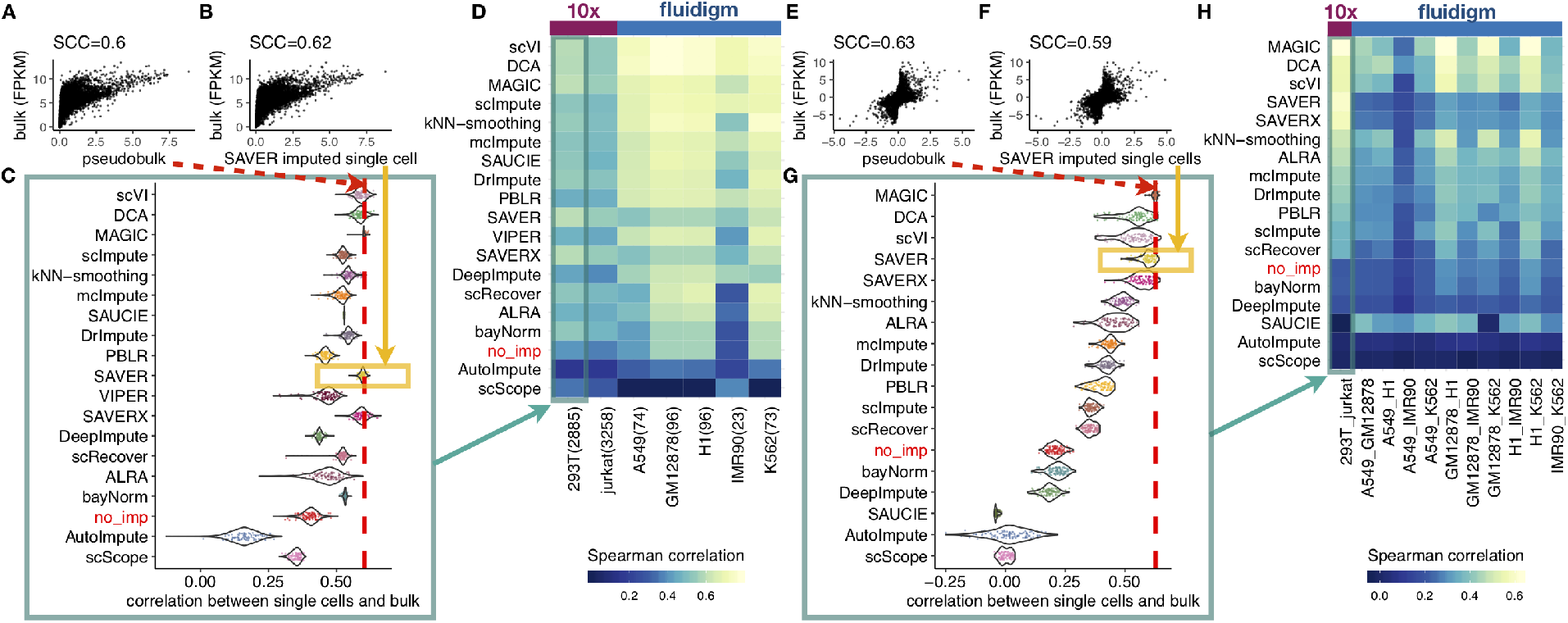
Similarity between bulk and imputed single-cell expression data in cell lines. **(A)** For the 293T cell line, a scatter plot of the `scran` normalized^37^ log2-transformed scRNA-seq cell profiles (*N* = 2885) averaged across all cells (’pseudobulk’) compared to a bulk RNA-seq profile with the Spearman’s correlation coefficient (SCC). **(B)** For each cell, we also calculated the SCC between an imputed cell’s profile (e.g. using SAVER) and the bulk RNA-seq profile. **(C)** Distribution of correlations between bulk profiles and single-cell profiles (imputed or not imputed – i.e. *no_imp*) across all cells in a the 293T cell line dataset. The red dotted line represents the estimated SCC 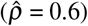 shown in (A). The methods are ordered in the same order as Figure (D) for comparison. **(D)** A heatmap of the median correlation for each imputation method and each dataset across two experimental platforms (two datasets from the 10X Genomics platform and five datasets from the Fluidigm platform with the number of cells in each dataset in parentheses). The rows are sorted by first averaging the median correlations across datasets within each platform and then averaging across platforms. **(E-H)** Similar to (A-D) except, for any two cell types, the SCC is calculated comparing the difference in two bulk cell type profiles compared to two scRNA-seq cell type profiles.

In the first evaluation, we directly compared imputed scRNA-seq profiles from cell lines to a bulk RNA-seq profile from the same cell lines (Figure 2A-D). The test data include 10x Genomics UMI-based scRNA-seq data for 293T and Jurkat cell lines (*10x_293T_jurkat*) and Fluidigm C1 plate-based scRNA-seq read count data for five ENCODE cell lines (*ENCODE_fluidigm_5cl*). Using the rank-based Spearman correlation coefficient (SCC)^41^ between the imputed scRNA-seq profile and bulk profile, the majority of imputation methods (16 out of 18) were found to improve the correlation compared to not imputing. Methods such as SAVER and SAVER-X (without pre-training) performed well with 10x Genomics UMI count data, but the method’s performance gain was not as pronounced with read count data from the plate-based Fluidigm platform, as expected^20, 21^. Other methods such as scVI, DCA and MAGIC performed well in both settings.

In the second evaluation, we assessed an imputation method’s ability to preserve the difference on the log scale between two cell type profiles (i.e. two cell lines) by comparing the difference in two single-cell cell type profiles to the difference in two bulk cell type profiles (Figure 2E-H). Compared to Figure 2A-D, the majority though a smaller set of imputation methods (13 out of 18) preserved the cell type difference better than no imputation. The imputation methods MAGIC, scVI, and DCA resulted in the highest correlation using both UMI and non-UMI plate-based protocols, but SAVER and SAVER-X (without pretraining) resulted in the highest correlation using UMI count data. Finally, in both Figures 2D and 2H, the performance of some imputation methods was found to be affected by the number of cells in the dataset with a smaller number of observations resulting in smaller correlation.

### 2.2 Impact of scRNA-seq imputation on identifying differentially expressed genes

Next, we evaluated the impact of imputation on the downstream analysis of identifying differentially expressed genes (DEGs). We intentionally designed our evaluation to primarily rely on empirical analyses of real data in order to preserve gene-gene correlations. In these empirical analyses, the ground truth was not completely known. Thus, our evaluation could not explicitly calculate sensitivity and specificity as previously studies^14, 15^. However, we preferred empirical evaluation over simulation. This is because some imputation methods model the expression levels for one gene based on the expression levels of other genes, such as with SAVER and SAVER-X, simulation or spike-in studies in which ground truth is known but the gene-gene correlation is disrupted would unfairly disfavor such methods. Also, modeling the complex gene-gene correlation in real data via simulation and spike-in studies is difficult.

In the first analysis, we performed a differential expression enrichment analysis (Figure 3A). We treated genes identified as differentially expressed in bulk RNA-seq data as a “gold standard” similar to previous studies^42^. We calculated the overlap of DEGs between the bulk data and DEGs identified from scRNA-seq data between the same two cell types (Figure S2A-G) using two methods for single-cell differential expression, namely MAST^43^ and Wilcoxon rank-sum test^44^ (abbreviated as Wilcoxon) and compared the performance (Figure 3B-D). Different from the differential analysis in Figure 2E-H which only considers the log-fold change (LFC) between cell types without taking into account cell variability, the analysis in Figure 3B-D additionally considers cell variability and thus the uncertainty of LFC in order to rank genes, similar to what one would do in hypothesis testing.

**Figure 3.**
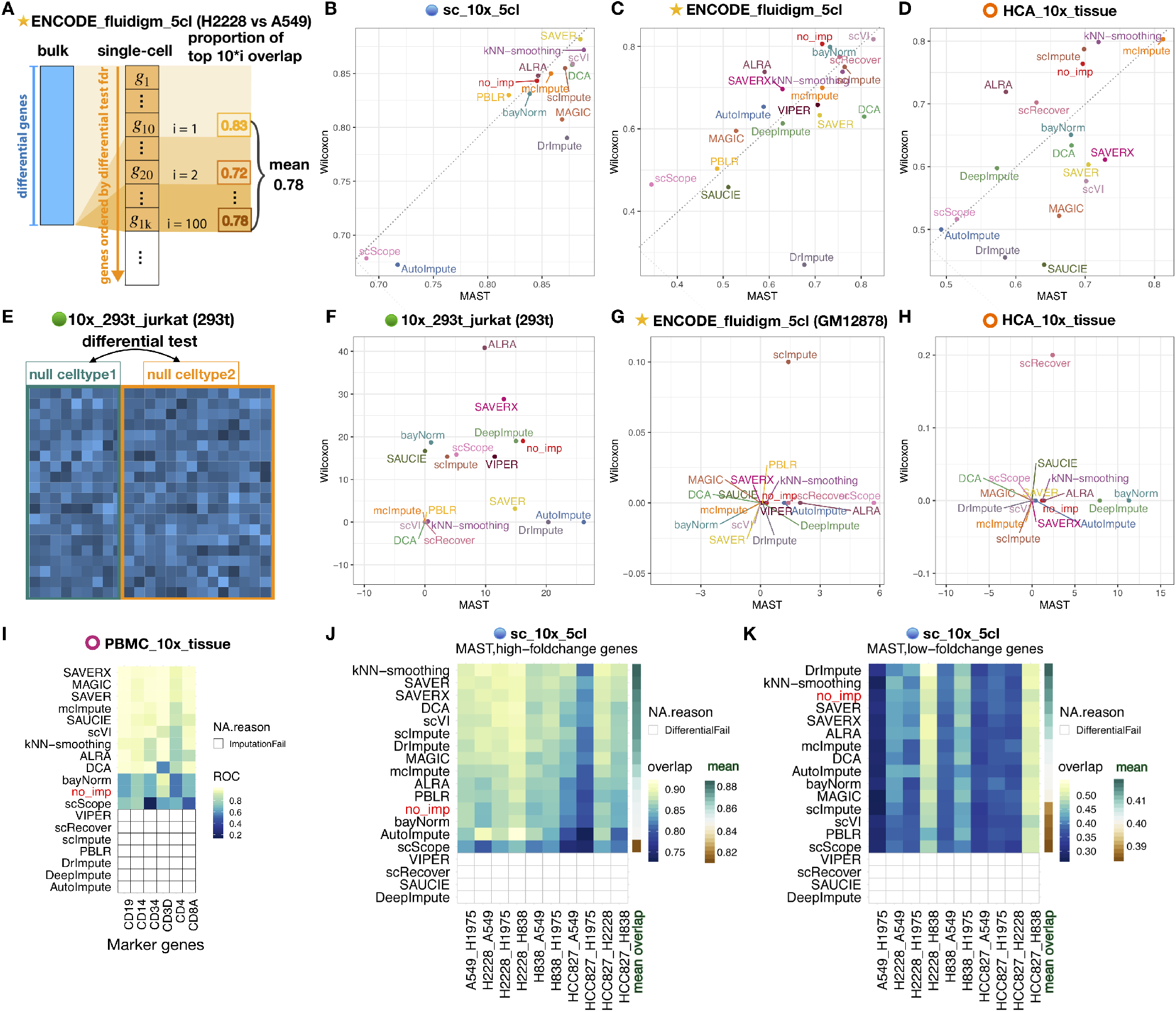
Impact of imputation methods on differential expression analysis. For each imputation method, we performed three gene-level analyses. **(A)** Schematic view of evaluating differentially expressed genes (DEGs) using the overlap between bulk RNA-seq and scRNA-seq. **(B-D)** Proportion of overlap between bulk and single-cell DEGs identified using either MAST (x-axis) or Wilcoxon rank-sum test (y-axis). Note that *‘cl’* in the names of datasets means ‘cellline’. **(E)** Schematic of a null differential expression analysis. **(F-H)** Number of false positive DEGs identified using MAST (x-axis) or Wilcoxon rank-sum test (y-axis). **(I)** Heatmap of area under a receiver operating characteristic (ROC) curve values when using the expression level of a marker gene (e.g. CD19) to predict a cell type (e.g. B cell or not) using UMI-based sorted PBMC cell types. For some imputation methods, no imputed values were returned. They are denoted as “ImputationFail”. **(J-K)** Using a UMI-based scRNA-seq dataset from cell lines (*sc_10x_5cl*), a heatmap showing the percentage of the overlap between bulk and single-cell DEGs identified using MAST stratified by genes with high (top 10%) or low (bottom 10%) log-fold changes. The color bar on the last column shows the mean overlap across all comparison for each method. If MAST failed to identify DEGs from the imputed profiles of any method in any dataset, we denoted it as “DifferentialFail”.

Using UMI-based 10x Genomics scRNA-seq data from five cell lines (*sc_10x_5cl*), we found that 10 and 8 out of the 18 methods outperformed no imputation when using MAST and Wilcoxon, respectively. Among them, kNN-smoothing, SAVER and SAVER-X (without pretraining) had the highest overlap of single-cell DEGs with bulk DEGs using MAST, but SAVER, kNN-smoothing and scVI performed the best when using Wilcoxon (Figure S2A-C). MAGIC had worse performance in this hypothesis testing setting (Figure S2B) compared to only calculating effect sizes (log-fold changes) without considering variance (Figure 2H). Further investigation shows that the estimated gene-specific standard deviations from the MAGIC imputed values were much smaller compared to the estimated standard deviations from other imputation methods (Figure S3). In turn, the estimated standard errors for the log-fold changes using MAGIC were small, leading to a wide test-statistic distribution and a *p*-value distribution skewed toward small *p*-values (Figure S4). These suggest that MAGIC could have inaccurately estimated cell variability which reduced its gene ranking performance.

On the plate-based read count data from five cell lines from ENCODE (*ENCODE_fluidigm_5cl*), scVI performed best using either MAST or Wilcoxon (Figure S2D-E). However, only 7 and 1 out of the 18 methods increased the overlap of DEGs compared to no imputation when using MAST and Wilcoxon, respectively.

We also compared methods using the UMI-based scRNA-seq bone marrow tissue data (*HCA_10x_tissue*) in 10x platform. Because cells in this dataset were unsorted, we first computationally labeled cell types using bulk RNA-seq data (Section 5.7). In this analysis, mcImpute and kNN-smoothing were in the top three either using MAST or Wilcoxon (Figure S2F-G). Again, the majority of imputation methods did not improve over no imputation. Only 6/18 and 3/18 methods outperformed no imputation when using MAST and Wilcoxon respectively.

Combining these results, broadly we found smoothing-based (e.g. kNN-smoothing) and data reconstruction methods (e.g. scVI) had the highest overlap with bulk DEGs across methods for differential expression and experimental protocols, which is consistent with previous results^14^. However, the model-based methods SAVER and SAVER-X performed well with UMI-based datasets.

In the second analysis, we performed a null differential expression analysis using cells from a homogeneous population where we expect no DEGs after correction for multiple testing. We selected a cell line dataset with a large number of cells, namely the H293T cell line from the *10x_293t_jurkat* dataset. We randomly split cells from the same cell type into two groups (Figure 3E) with varying group sizes ranging from *N* = 10 to 1000 cells per group (Figure S2H-N), imputed the expression values and identified DEGs (Figure 3F-H). Broadly, we found no consistency in which types of imputation methods identified false positive DEGs (Figure S2I-N) as there were examples in all types of imputation methods (model-based, smoothing-based and data reconstruction based) that reported false positive DEGs. However, one factor associated with these false positives was found to be an imbalanced number of observations in each group when identifying DEGs, using both MAST and Wilcoxon (Figure S2I-J).

In the third analysis, we asked if the imputed expression of known cell-type-specific marker genes can correctly predict cell type. Using a UMI-based and FACS sorted peripheral blood mononuclear cell (PBMC) dataset^3^ (*PBMC_10x_tissue*) and known PBMC marker genes highly expressed in purified PBMC cell types (CD19 for B cells; CD14 for monocytes; CD34 for CD34+ cells; CD3D for CD4 T helper cells, cytotoxic T cells, memory T cells, naive cytotoxic T cells, naive T cells, regulatory T cells; CD8A for cytotoxic T cells and naive cytotoxic T cells), we evaluated the performance of predicting a cell type (e.g. B cell) based on the expression of a marker gene (e.g. CD19) (Figure 3I). We estimated the area under the ROC (AUROC) curve where the expression of the marker gene (e.g. CD19 expression) is the predictor and the true cell type (e.g. B cell or not) is the label (for more details see Section 5.5). Among the methods that returned imputed values, most methods (10/11) produced higher AUROC compared to no imputation. Specifically, SAVER-X, MAGIC, and SAVER were the top three methods with this UMI-based scRNA-seq data.

We further evaluated the impact of the magnitude of differential expression on an imputation method, which was previously shown to be important for the performance of imputation methods in the context of DEG analysis^14^. Using the UMI-based scRNA-seq dataset with five cell lines (*sc_10x_5cl*), we compared cell lines pairwise and stratified genes into high and low LFC using the bulk RNA-seq data, where high (or low) LFC genes were defined as the top 10% (or bottom 10%) LFC. Interestingly, when the LFC magnitude was large, the overlap between the bulk and single-cell DEGs for most imputation methods (11/14) was higher compared to no imputation (Figure 3J). However, when the LFC was small, only 2/14 methods had larger overlap between the bulk and single-cell DEGs compared to no imputation, suggesting that most imputation methods may have smoothed small differential signals away (Figure 3K).

### 2.3 Impact of scRNA-seq imputation on unsupervised clustering

Unsupervised clustering is another common downstream analysis with scRNA-seq data to empirically define groups of cells with similar expression profiles^40^. Here, we assessed the impact of the 18 imputation methods on unsupervised clustering, specifically using *k*-means^45^ and Louvain clustering^46^. Clustering was performed on the top principal components, but we also evaluated the latent spaces directly provided by scVI, scScope and SAUCIE, which increased the total number of methods to 21. We used four metrics to assess the clustering performance: the median Silhouette index, adjusted Rand index (ARI)^47^, entropy of cluster accuracy (ECA or *H_accuracy_*), and entropy of cluster purity (ECP or *H_purity_*). The last three were also used by and described in Tian et al. (2019)^18^. The Silhouette index measures consistency within clusters (or how similar an observation is to its own cluster compared to other clusters). The last three metrics assess the similarity of predicted cluster labels to a known ground truth, and they have been shown to have good correlation with each other^18^. The primary difference between the last two is that ECA measures the diversity (or accuracy) of the true group label within each cluster assigned by the clustering method, while ECP measures the diversity (or purity) of the predicted cluster labels within each of the true groups. We scaled ARI to range between 0 and 1, and Silhouette index ranges between −1 and 1 – in both cases a higher score represents a better performance, but ECA and ECP range between 0 and a number larger than one with a lower score representing a better performance.

We applied each imputation method to seven datasets from CellBench^18^ (Table S1, Section 5.1), a data compendium consisting of both UMI-based 10x Genomics, Drop-seq and plate-based scRNA-seq data for benchmarking analysis methods, and then applied *k*-means clustering (Figure 4, Figure S5). A similar evaluation using the CellBench data was performed in Tian et al. (2019)^18^ who evaluated three scRNA-seq imputation methods (kNN-smoothing^27^, DrImpute^25^, and SAVER^20^) with five unsupervised clustering methods. Here, we expanded their analysis to include 21 scRNA-seq imputation methods with our goal of prioritizing the evaluation of the imputation methods. Broadly, using *k*-means clustering only 5 of 21 methods (MAGIC, mcImpute, SAVER-X, SAVER and scImpute) improved clustering results compared to no imputation in this data (Figure 4A). Overall, using latent spaces for the methods scVI (e.g. observations sampled from the posterior distribution using scVI), scScope and SAUCIE was better than using their imputed expression values. We illustrate how individual cells cluster along the first two principal components using an example dataset from CellBench (*sc_celseq2_5cl_p1*) with no imputation and with imputation using MAGIC (Figure 4B). Using Louvain clustering yielded similar results (Figure 4C, Figure S6) although the top methods were slightly different (MAGIC, SAVER-X, SAVER, ALRA, bayNorm).

**Figure 4.**
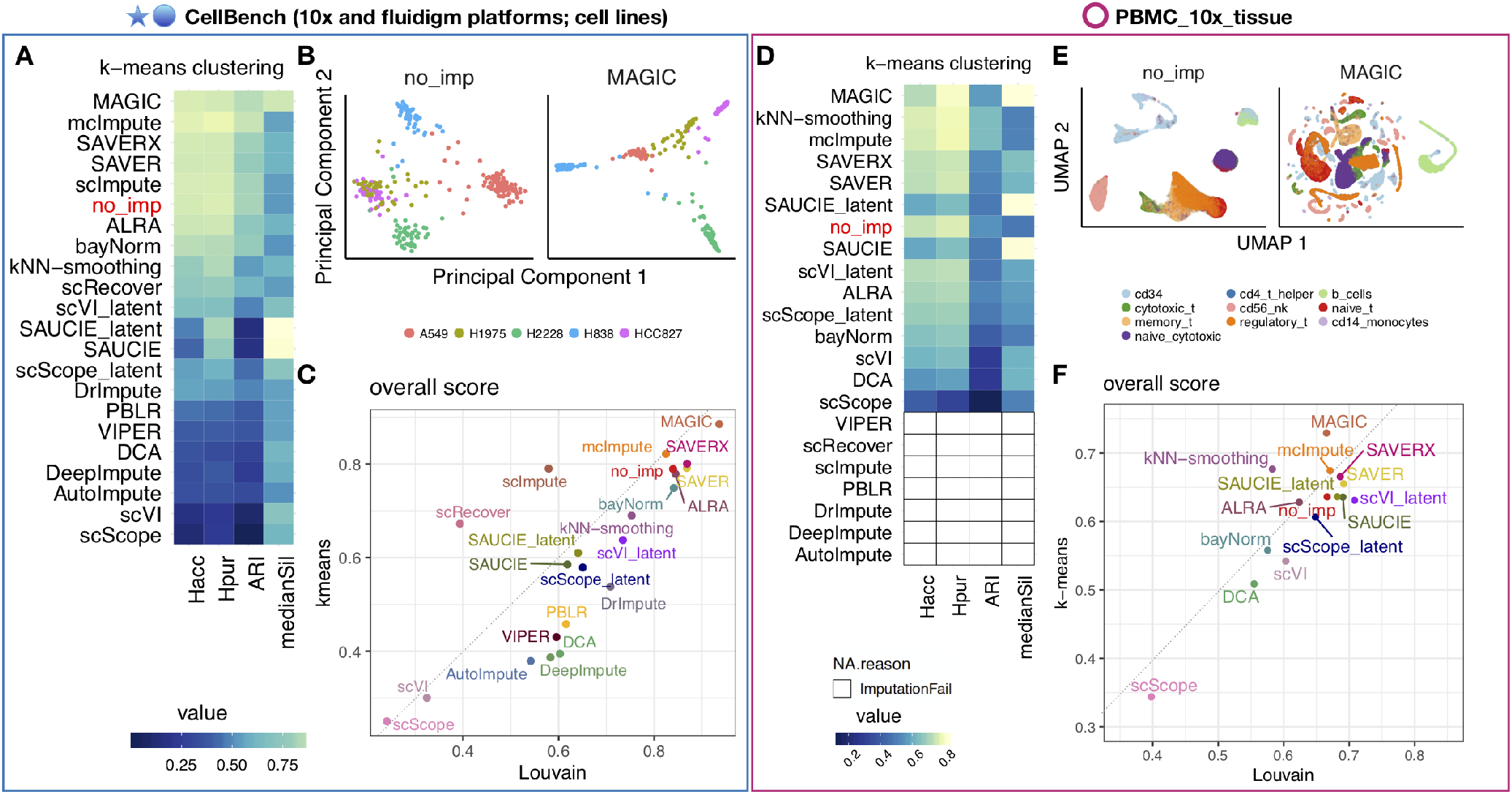
Impact of imputation methods on unsupervised clustering analysis. **(A)** Heatmap of four performance metrics – entropy of cluster accuracy (*H_acc_*), entropy of cluster purity (*H_pur_*), adjusted Rand index (ARI), and median Silhouette index – averaged across seven datasets from CellBench^18^. Each metric shows the average performance across 7 datasets in CellBench. To compare imputation methods across metrics, the metrics were re-scaled to between 0 and 1 and the order of *H_acc_* and *H_pur_* were flipped to where a higher standardized score translates to better performance. Imputation methods (rows) are ranked by the average performance across all four metrics. **(B)** Dimension reduction results after applying PCA to the *sc_celseq2_5cl_p1* data with no imputation (left) and with imputation using MAGIC (right). The colors are the true group labels. **(C)** Overall score (or average of the four performance metrics) for Louvain clustering (x-axis) and *k*-means clustering (y-axis). **(D-F)** Same as (A-C) except using the scRNA-seq dataset of ten sorted peripheral blood mononuclear cell (PBMC) cell types from 10x Genomics^3^ (*PBMC_10x_tissue* dataset). White areas with black outline in (D) indicate that the imputation methods did not return output after 72 hours. Also, Figure (E) uses UMAP components^48^ instead of principal components.

We further compared imputation methods using the scRNA-seq dataset of ten sorted PBMC cell types from 10x Genomics^3^ (*PBMC_10x_tissue*). We applied both *k*-means clustering and Louvain clustering. For the imputation methods that successfully returned imputed values, 6 methods including MAGIC, kNN-smoothing, SAVER-X, SAVER and SAUCIE_latent (the latent space output by SAUCIE) outperformed no imputation (Figure 4D). In Figure 4E, we show how the sorted PBMC cells cluster along the UMAP components^48^ with no imputation and with imputation using MAGIC. Again, the results between *k*-means and Louvain clustering were similar (Figure 4F, Figure S7) though the top methods when using Louvain clustering were slightly different (scVI_latent, SAVER, SAUCIE, SAVER-X, SAUCIE_latent, mcImpute).

### 2.4 Impact of scRNA-seq imputation on inferring pseudotemporal trajectories

We also evaluated the impact of imputation methods on inferring cells’ pseudotemporal trajectories. In contrast to Section 2.3 which used data with distinct cell types to evaluate clustering, here we used datasets in which cells had a continuum of transcriptomic profiles (e.g. cell differentiation) to assess if imputation methods could recover continuous biological processes. Analogous to the section above, we applied methods to infer trajectories on the top principal components. We also included the latent spaces directly provided by scVI, scScope and SAUCIE.

First, we applied imputation methods to six RNA mixture and cell mixture datasets from CellBench^18^ followed by using two trajectory analysis methods, Monocle 2^50^ and TSCAN^49^. In these data, the true trajectories of cells were known. They were used to evaluate the impact of imputation methods on the ability to infer trajectories. A similar evaluation using CellBench data was performed by Tian et al. (2019)^18^ who evaluated three scRNA-seq imputation methods (kNN-smoothing^27^, DrImpute^25^, and SAVER^20^) with five trajectory inference methods. Here, we expanded their analysis to include 21 scRNA-seq imputation methods. Our primary goal is to evaluate the imputation methods, but we do include two trajectory inference methods. The performance metrics used in this analysis were (1) the Pearson *correlation* between cells’ rank order along the inferred trajectory and their rank order along the true trajectory, and (2) the proportion of cells for which the inferred branch *overlapped* (i.e., was consistent) with the branch in the true trajectory. Both of these metrics have been previously described in and used to evaluate inferred cell trajectories^18^.

Using the CellBench data, we found the imputation methods kNN-smoothing, SAVER, and ALRA led to both increased correlation (Figure 5A) and overlap (Figure 5B) compared to no imputation using the TSCAN trajectory inference. Using Monocle 2 trajectory inference, SAVER, kNN-smoothing, mcImpute, and the latent spaces from SAUCE (SAUCE_latent) increased both the correlation (Figure S8A) and overlap (Figure S8B) compared to no imputation. This confirms the variability in imputation methods’ performance (depending on trajectory inference method), in particular for the overlap, that was previously shown^18^ (Figure 5C-D). Finally, analogous to results shown in Section 2.3, for imputation methods that return latent spaces, using the latent spaces generally led to better performance than using the imputed expression values.

**Figure 5.**
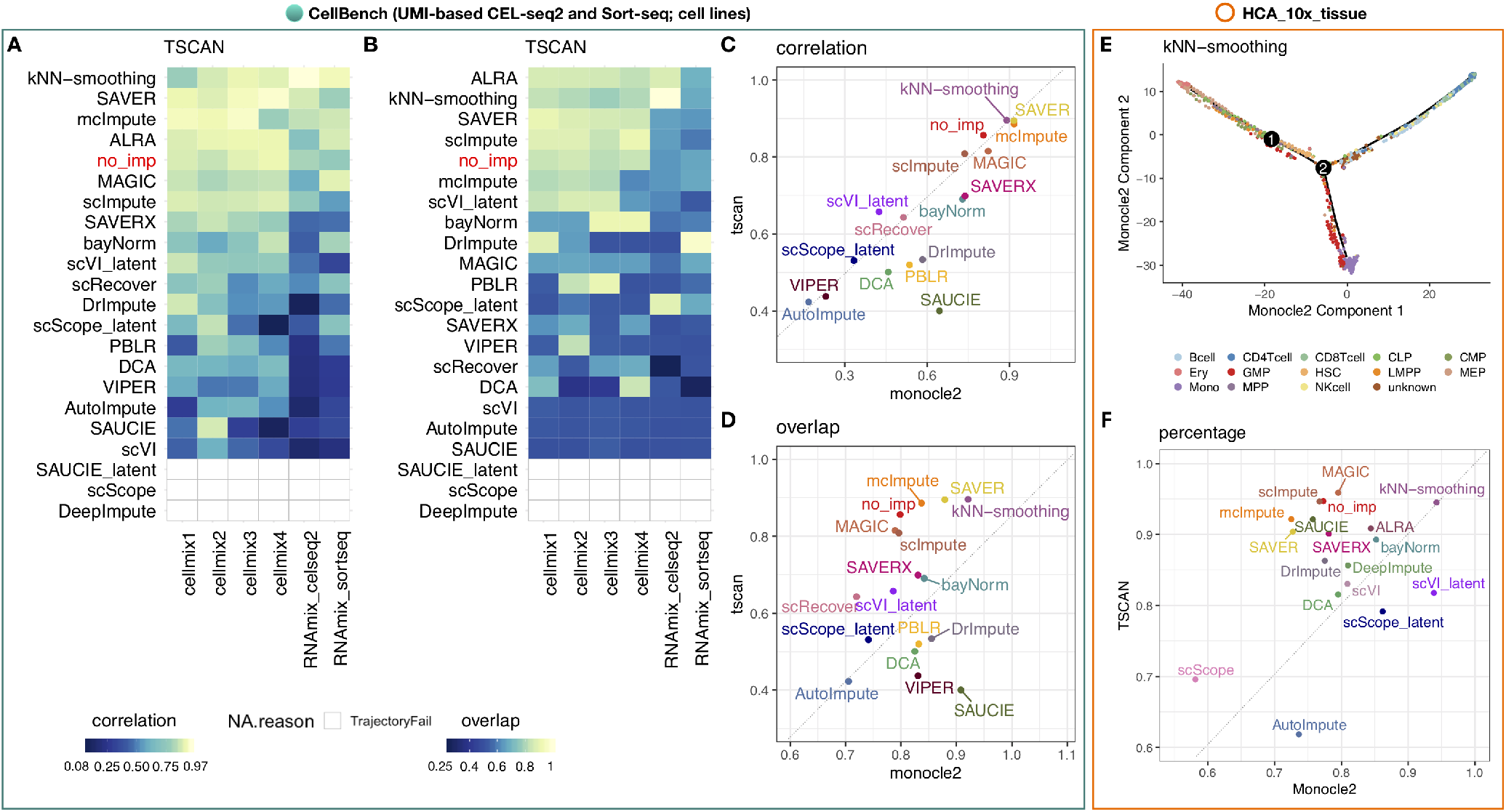
Impact of imputation methods on inferred trajectories for pseudotime analysis. **(A)** Heatmap showing the Pearson correlation coefficients (PCC), denoted as *correlation*, between the ranks of the inferred trajectory using TSCAN^49^ and the rank order of the cells where we know the true trajectory (or ordering) of the cells, using the six RNA mixture and cell mixture datasets from CellBench^18^. White areas with grey outline in (A) and (B) indicate that TSCAN failed to infer trajectories from the imputed profiles. **(B)** Heatmap of the proportion of cells on the inferred trajectories from TSCAN that correctly *overlap* with the cells on the branch where we know the true trajectory of the cells using the same data as (A). **(C)** The comparison of the correlation and **(D)** overlap averaged across datasets using Monocle 2^50^ (x-axis) and TSCAN^49^ (y-axis) as the trajectory reconstruction method. **(E)** An inferred trajectory from Monocle 2 using *N*=6941 bone marrow cells from the *HCA_10x_tissue* that were imputed using kNN-smoothing^27^. Colors represent computationally defined cell types using bulk RNA-seq data (see Section 5.7 for details). **(F)** Here, we compare the estimated pseudotime to the level of differentiation for each pair of bone marrow cells. For instance, for the pair of cells A and B, if cell A is a hematopoietic stem cell (HSC) (we assign it as differentiation level 1), cell B is a Multi-Potent Progenitor (MPP) cell (we assign it as differentiation level 2), and the inferred pseudotime for cell A and B is *t_A_* and *t_B_* where *t_A_* < *t_B_*, then we call it “correctly ordered”. The *percentage* of the correctly ordered cells averaged across all possible cell pairs serves as the assessment measure considering both Monocle 2 (x-axis) and TSCAN (y-axis).

Next, we evaluated the performance of the imputation methods using bone marrow cells from the *HCA_10x_tissue*^11^. Because these are not cell lines, we first computationally labeled cell types using bulk RNA-seq data (see Section 5.7 for details). We used the schematic of hematopoietic stem cell (HSC) differentiation shown in Figure 1A in Buenrostro *et al.*^51^ to computationally label cell types. The bone marrow contains hematopoietic stem cells (HSCs) differentiating into three major lineages: lymphoid, erythroid, and myeloid cells (Figure 5E). Here, we compare the estimated pseudotime to the level of differentiation for each pair of bone marrow cells. For instance, for the pair of cells A and B, if cell A is a HSC cell (we assign it as differentiation level 1), cell B is a multipotent progenitor (MPP) cell (we assign it as differentiation level 2), and the inferred pseudotime for cell A and B is *t_A_* and *t_B_* where *t_A_* < *t_B_*, then we call it “correctly ordered”. The *percentage* of the correctly ordered cells averaged across all possible cell pairs serves as the performance metric.

Similar to using the CellBench data, kNN-smoothing most consistently outperformed other imputation methods using this heterogeneous tissue data using either TSCAN or Monocle 2. However there was variability in the performance of other imputation methods depending on the trajectory analysis method used (Figure 5F), as previously reported^18^. Figure 5F shows the percentage of correctly ordered cell pairs in the bone marrow dataset using Monocle 2 (x-axis) and TSCAN (y-axis) as the pseudotime reconstruction methods. We found kNN-smoothing and MAGIC slightly performed better than using no imputation. However, the majority (all but two) of methods reported 78% correctly ordered cells in this data using either TSCAN or Monocle 2.

## 3 Discussion and Conclusions

We have presented a systematic benchmark evaluation comprehensively comparing 18 scRNA-seq imputation methods. We evaluated the performance of the imputation methods on their ability to recover similarity between imputed single-cell profiles and bulk profiles in a homogeneous population of cells, and the impact of the imputation methods on three downstream analyses: differential expression analysis, unsupervised clustering analysis, and pseudotime inference. We conclude by summarizing our results in Figure 6 which ranked methods based on their average performance. Computational time, memory usage, and scalability were not used to rank the methods. They were assessed separately using four datasets of 10^3^, 5 × 10^3^, 5 × 10^4^ and 10^5^ cells, respectively. Specifically, we used (1) computation time (in seconds), (2) memory (in maximum resident set size of all tasks in job, i.e. MaxRSS, returned from `sacct`), and (3) scalability – regression coefficient in the linear model where the computation time is fitted against the number of cells on the log_10_-scale. Figure S9 shows the comparison of time, memory and scalability. Based on our evaluation, we provide a set of recommendations below for users and investigators to navigate the current space of scRNA-seq imputation methods.

**Figure 6.**
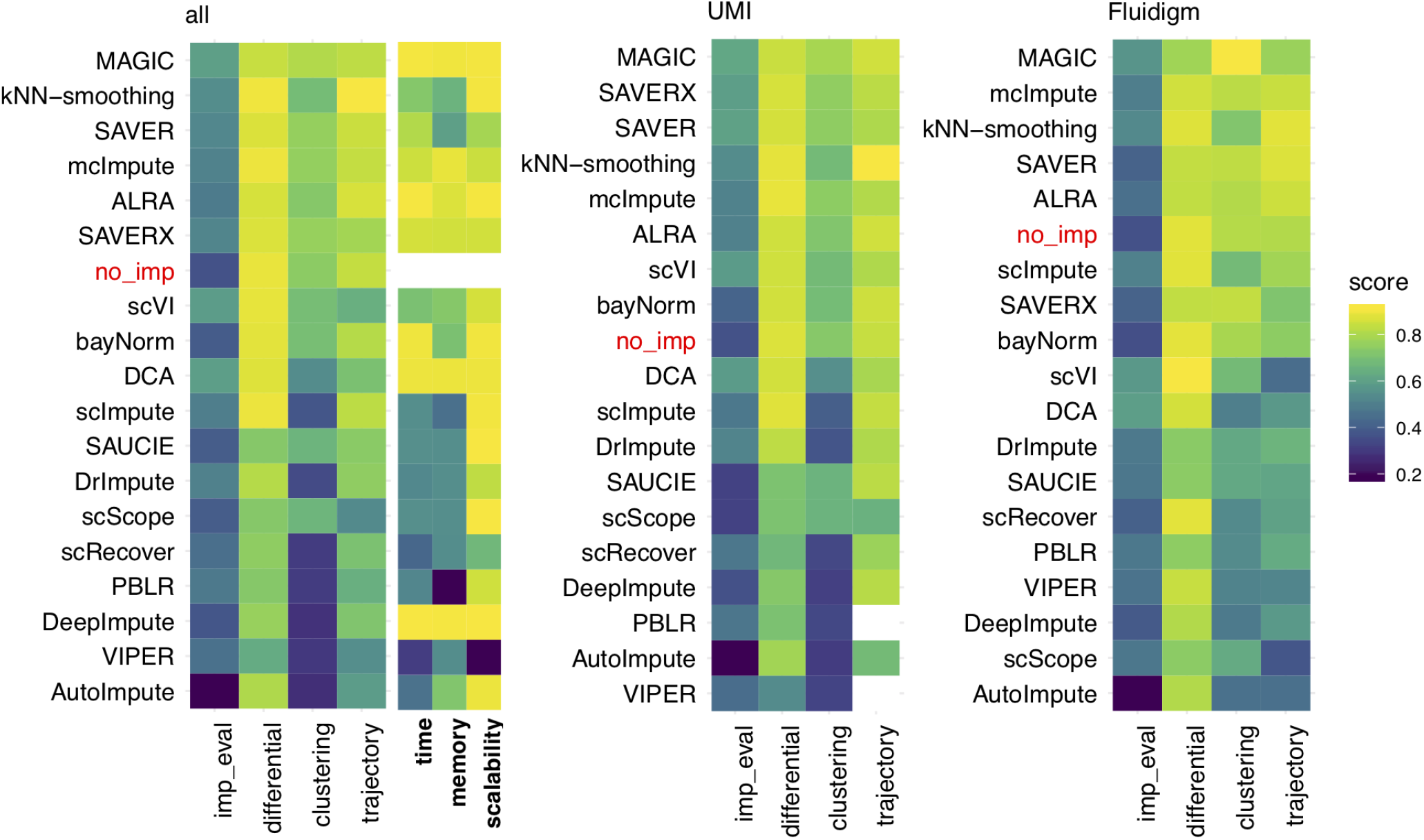
Overall summary of results evaluating imputation methods for scRNA-seq data. Methods are ranked by performance metrics (scaled to apply the same color scale) averaged across four categories: evaluation of similarity between imputed single-cell profiles and bulk profiles in a homogeneous population of cells (imp_eval), differential expression enrichment analysis and null differential analysis (differential), unsupervised clustering and inferring trajectories for pseudotime analysis. A higher score represents a better performance. Results for using UMI or non-UMI (e.g. in this case Fluidigm) methods are also shown.

Of the methods considered, MAGIC, kNN-smoothing and SAVER were found to outperform the other methods most consistently (Figure 6). However, the performance of methods varied across evaluation criteria, experimental protocols, datasets, and downstream analyses. For example, scVI was one of the top performers in terms of the similarity between the imputed single-cell and bulk expression profiles (Figure 2), but it did not perform among the top in clustering and trajectory analysis (Figures 4,5). SAVER-X performed consistently well using UMI-based method, but less well with non-UMI based methods (Figure 6). While MAGIC was one of the best performer overall (Figure 6), it performed worse than many other methods when identifying differentially expressed genes in hypothesis testing type settings that take into account cell variability (Figure 3).

In addition, we found that while some imputation methods improve detecting differentially expressed genes or discovering marker genes, they also can introduce false positive signals, often driven by imbalanced sizes of groups. The magnitude of effect sizes (i.e. log-fold change) plays a role in the performance of the imputation methods: most imputation methods strengthen large effect sizes compared to no imputation. However, if the original expression difference is small, then most imputation methods smooth the small differential signal away and do not show clear advantage over no imputation (Figure 3J,K).

One important observation is that, while the majority of imputation methods outperformed no imputation in recovering bulk expression (16/18 methods) and log fold changes of individual genes between cell types without considering cell variability within each cell type (13/18 methods) (Figure 2), much fewer methods performed better than no imputation for identifying differentially expressed genes after considering cell variability (1-10/18 depending on the test scenario), clustering cells (5-6/21 methods) or inferring pseudotemporal trajectories (4-11/21 methods). Thus, the current imputation methods as a whole seem to be most effective for providing a point estimate of the activity of individual genes, and they become less effective when coupled with various downstream analysis tasks. For differential expression analysis, the decreased effectiveness is likely due to inaccurate cell variance characterization after imputation. For clustering and trajectory analysis, the reduced effectiveness is likely because these two analyses attempt to analyze cell-to-cell relationship rather than individual genes. Cell clustering and trajectory analysis are usually conducted by embedding the high-dimensional expression vector of each cell into a relatively low-dimensional space. Each dimension in the low-dimensional space combines information from many genes, which increases signal-to-noise ratio by diluting technical noise such as observed zeros due to technical variation, even without imputation. Thus, the recovery of cell-to-cell relationship can be influenced less by imputation. By contrast, the measurements of individual genes contain high level of technical noises which can be greatly mitigated by imputation by borrowing information from other genes or cells. Thus, imputation could be more helpful for analyzing individual genes rather than cell-to-cell relationship. An open question to be investigated in the future is whether the improvement on the various downstream analysis tasks by imputation has already reached its upper limit, and if not, how to design new imputation methods to further improve the analysis of cell-to-cell relationship or differential expression that takes into account cell variability.

In terms of computation, MAGIC, DCA and DeepImpute are among the most efficient methods. *k*NN-smoothing, ALRA, bayNorm, scImpute, SAUCIE, scScope exhibit high scalability. SAVER-X, SAVER and SAVER-X are intermediary while the remaining methods do not scale well for large datasets (Figure S9).

Our comparison is subject to several limitations. Firstly, the imputation methods were mostly compared with default parameters which may not achieve optimal performance across all datasets. Our work could be further improved with the use of methods such as molecular cross-validation (MCV)^52^. Another limitation is we used 72 hours as the time limit for convergence for imputation methods, which does not guarantee algorithmic convergence for some methods. In our evaluation of imputation methods on inferring pseudotime with trajectory analysis methods, the cell types of the tissue *HCA_10x_tissue* cells were computationally annotated. However, our benchmark evaluation will benefit many current researches using scRNA-seq data as it highlights the advantages and disadvantages of existing imputation methods, and that the performance of an imputation method depends on many external factors, such as experimental protocols and analyses usage. It also provides an evaluation standard for new imputation methods.

## 4 Back Matter

### 4.1 Author contributions

All authors conceptualized and contributed to the main aims of the project. WH and ZJ performed the benchmark data analyses. All authors wrote and approved the final manuscript.

## 5 Methods

All methods were evaluated with default parameters, with the exception of the deep-learning-based methods for which the maximum epoch time was set as 400. We used 72 hours as the time limit for convergence for the imputation methods. This did not guarantee algorithmic convergence for some methods. For a description of the data, see Section 5.1 and Table S1. For complete details on the methods used, the input, the output, pre-processing steps, the programming language used, version number and link to software, see Table S2.

### 5.1 Data

#### 5.1.1 Single cells and bulk samples from cell lines

1. *Jurkat* cell lines

- *10x_293t_jurkat* (293T cells): *N*=3258 cells measured using UMIs and the droplet-based protocol from the 10x Genomics Chromium^3^ (link to data)
- *N*=2 bulk RNA-seq sample (GSE129240^53^)
2. *HEK293T* cell lines

- *10x_293t_jurkat* (jurkat cells): *N*=2885 cells measured using UMIs and the droplet-based protocol from the 10x Genomics Chromium^3^ (link to data)
- *N*=2 bulk RNA-seq sample (GSE129240^53^)
3. Five ENCODE (*A549*, *GM12878*, *h1-hESC*, *IMR90*, *K562*) cell lines

- *ENCODE_fluidigm_5cl*: The five cell lines correspond to five cell types and contain a total of *N*=362 cells all from GSE81861. They were sequenced with the SMARTer full-length method^17^ using the Fluidigm C1 protocol^54^. The cell types include *N*=74 A549 cells, *N*=96 GM12878 cells (batch 2), *N*=96 H1-hESC cells (batch 1) referred to as *H1* in the manuscript, *N*=23 IMR90 cells, and *N*=73 K562 cells.
- The bulk RNA-seq samples from ENCODE^55^: *N*=7 A549 samples, *N*=11 GM12878 samples, *N*=8 H1-hESC samples, *N*=5 IMR90 samples, and *N*=27 K562 samples
4. Single cells and pseudo cells from the CellBench^18^ scRNA-seq benchmarking dataset

- All UMI-based data from five cell lines (HCC827, H1975, H2228, H838, A549) in the CellBench^18^ benchmarking dataset (except for *cellmix5*, a population control) using the CEL-seq2 protocol (*sc_celseq2*, *sc_celseq2_5cl_p1*, *sc_celseq2_5cl_p2*, *sc_celseq2_5cl_p3*, *cellmix1*, *cellmix2*, *cellmix3*, *cellmix4*, *RNAmix_celseq2*), Drop-seq Dolomite protocol (*sc_dropseq*), the Sort-seq protocol (*RNAmix_sortseq*), and 10x Chromium Genomics protocol (*sc_10x*, *sc_10x_5cl*)). For a description of the experimental design, GEO accession numbers, protocol parameters, see the sc_mixology GitHub repo and Table S1.
- *N* = 10 bulk RNA-seq samples from GSE86337^56^ (each of the five cell lines have two replicates)

#### 5.1.2 Single cells and bulk samples from tissues

1. Bone marrow tissue from the Human Cell Atlas^11^ (HCA)

- *HCA_10x_tissue*: *N*=6939 bone marrow cells from sample MantonBM6 measured using 10x Genomics link to data
- *N*=49 bulk RNA-seq samples from 13 cell types (B cell, CD4 T cell, CD8 T cell, CMP, GMP, HSC, MEP, Monocyte, MPP and NK cells each has 4 samples; and each of CLP, Erythroid and LMPP has 3 samples). (GSE74246)
2. Sorted peripheral blood mononuclear cell (PBMC) tissue from 10x Genomics (UMI)

- *PBMC_10x_tissue*: *N*=59620 sorted cells from 10 cell types (*N*=4033 B cells, *N*=498 CD14 monocyte cells, *N*=9162 CD34 cells, *N*=7046 CD4 T helper cells, *N*=7555 CD56 NK cells, *N*=7631 cytotoxic T cells, *N*=6969 memory T cells, *N*=5672 naïve cytotoxic cells, *N*=4569 naïve T cells, and *N*=6485 regulatory T cells) from the 10x Genomics^3^.

### 5.2 Data processing and imputation

#### 5.2.1 Single-cell RNA-seq data

We applied the same quality control (QC) criterion across all single-cell datasets. Cells with at least 500 detected genes were retained. ERCC spike-ins were removed. Genes expressed in at least 10% of cells in cell line data and 1% of cells in tissue samples were retained. Mitochondrial genes were removed. We applied these cell- and gene-filtering steps to all imputation methods and skipped each method’s own gene- and cell-filtering (if applicable) to keep the dimension of the imputed values (output) the same across imputation methods. Single-cell profiles from UMI-based *Jurkat* and *HEK293T* cell lines were combined into one UMI count matrix (*10x_293t_jurkat*) which was used as input to each imputation method. A similar procedure was applied to the five Fluidigm-based ENCODE cell lines (*ENCODE_fluidigm_5cl*). Data were normalized by the pooling normalization method^37^ implemented in the `scran`^38^ R/Bioconductor^39, 40^ package and log 2-transformed if any imputation method requires normalized counts or log-transformed normalized counts as input. We added post-processing steps for methods if required. For example, if a method did not normalize nor apply a log-transformation before or during imputation, we applied `scran` normalization and log 2-transformation to the output. If a method included normalization but does not log-transform the data during imputation, we applied log 2-transformation to the output.

#### 5.2.2 Bulk RNA-seq data

Read counts were downloaded from GEO (accession numbers GSE129240, GSE86337, and GSE74246). Each sample was normalized by library size and multiplied by a size factor 10^6^, or count per million (CPM), and then log 2-transformed with a pseudocount of 1, i.e. log 2(*CPM* + 1). For the ENCODE samples, the downloaded data were already normalized (transcripts per million or TPM) and log 2-transformed.

### 5.3 Null simulation

To generate Figure 1A and Figure S1, we used the 293T cells from the *10x_293t_jurkat* dataset with *G* = 18729 genes and *C* = 2885 cells. We applied gene-level quality control by only retaining genes that have expression at least 10% of cells, which kept *G*=7582 genes. Let *X_gi_* represent the observed scRNA-seq UMI counts for the *g^th^* gene where *g* ∈ (1,…, *G*) and the *i^th^* cell where *i* ∈ (1,…,*C*) from a given dataset. We estimated the mean expression level for each gene across the cells: 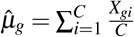. Then, for each gene, we simulated scRNA-seq UMI counts *Y_gj_* for the *g^th^* gene and the *j^th^* simulated cell from a Poisson distribution with the mean equal to the product of the estimated gene-level mean 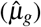 estimated from **X** and a cell-specific library size factor (*δ_j_*) randomly sampled from a uniform distribution between [*a, b*]:

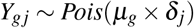

where *δ_j_* ~ *U* [*a, b*] for *j* ∈ (1,…,*C*′). For Figures 1A and S1, *C*′ = 1000 and the library size factors *δ_j_* ~ *U* [0.9, 1.1]. We passed the gene expression matrix to each imputation method. Data preprocessing and postprocessing steps followed Section 5.2. Principal component analysis was performed on genes with coefficient of variation (cv) greater than the median cv.

### 5.4 Evaluation of similarity between bulk and imputed scRNA-seq data

#### 5.4.1 Correlation of gene expression profiles between bulk and imputed scRNA-seq profiles

For a given scRNA-seq dataset, we averaged the `scran` normalized^37^ log 2-transformed scRNA-seq counts across cells (referred to as a ‘pseudobulk’) and calculated the Spearman’s rank correlation coefficient (SCC) (or *ρ*) between the pseudobulk and the bulk RNA-seq profile (averaged across replicate bulk samples) of the same cell type. Next, for each cell in a given scRNA-seq dataset, we calculated the SCC between the cell’s imputed scRNA-seq profile (e.g. using SAVER) and the averaged bulk RNA-seq profile. The median SCC across all cells was then used to evaluate the performance of an imputation method within a dataset. To rank methods across datasets, we first averaged the median SCCs across datasets within the same experimental protocol (e.g., UMI-based or plate-based) and then averaged these averages across protocol.

#### 5.4.2 Correlation between the bulk and imputed single-cell gene expression log fold changes

This analysis was similar to Section 5.4.1, but here the SCC was calculated by comparing bulk and single-cell differential expression. For a given pair of cell types, we first averaged the cell profiles from each cell type to form two ‘pseudobulk’ samples using `scran` normalized^37^ and log 2-transformed scRNA-seq profiles. The difference between the two pseudobulk profiles and the difference between the two bulk RNA-seq profiles were computed and the SCC between the two differential profiles was computed.

Next, we took one cell from each cell type and calculated the difference in the imputed scRNA-seq profile (e.g. using SAVER) between these two cells. The SCC between this difference and the difference between two bulk RNA-seq profiles from the same two cell types was then computed. This was repeated for all possible cell pairs. For a given pair of cell types (both within the same experimental protocol), the median SCC across all cell pairs was computed. To rank methods across datasets, we averaged the median SCCs across datasets within each protocol and then averaged these averages across protocols.

### 5.5 Evaluation of imputation to identify differentially expressed genes

#### 5.5.1 Ranking differentially expressed genes (DEGs)

For each of the three imputed scRNA-seq datasets (*sc_10x_5cl*, *ENCODE_fluidigm_5cl* and *HCA_10x_tissue*), we identified differentially expressed genes (DEGs) between all pairs of cell types. We considered two methods to identify DEGs from scRNA-seq: (1) MAST^43^ which models the data using a hurdle model and (2) Wilcoxon rank-sum test^44^. For bulk RNA-seq samples, DEGs were identified using the `limma`^57^ R/Bioconductor package. We corrected *p*-values for multiple testing using the Benjamini-Hochberg (BH) method^58^ (`p.adjust` function in the `stats` R package) to derive false discovery rate (FDR) (only when a method did not already correct for FDR) and identified genes with FDR smaller than *α* = 0.05. The bulk DEGs were treated as a “gold standard” similar to previous studies^42^. For the scRNA-seq data, The single-cell DEGs were ranked by *p*-values or the log-scaled expression fold change if there was a tie for *p*-values. For *i* from 1 to 100, we calculated the proportion of top 10 ∗ *i* single-cell DEGs that overlap with bulk DEGs. The average of these 100 proportions served as the performance metric.

#### 5.5.2 Null differential analysis

For each dataset in this analysis, we started with a homogeneous population of cells where we expect no DEGs after correction for multiple testing. We used the the 293T cells from the *10x_293t_jurkat* dataset (*N*=2885 cells), the GM12878 cell line from the *ENCODE_fluidigm_5cl* dataset (*N*=96 cells), and the cell type with the largest number of cells from the *HCA_10x_tissue* (*N*=193 monocytes). For each dataset, we randomly sampled cells into two groups with group size ranging from *N* = 10 to 1000 cells per group, imputed the expression values of these two groups together and identified DEGs using MAST^43^ and Wilcoxon rank-sum test^44^. We filtered for FDR smaller than *α* = 0.05.

#### 5.5.3 Predicting cell type using imputed expression of known PBMC marker genes

Using the sorted peripheral blood mononuclear cells (PBMCs)^3^(*PBMC_10x_tissue* dataset), we assessed the performance of an imputation method on recovering the expression level of known PBMC marker genes. The cell type-specific marker genes used in this analysis were the following: CD19 for B cells; CD14 for monocytes; CD34 for CD34+ cells; CD3D for CD4 T helper cells, cytotoxic T cells, memory T cells, naive cytotoxic T cells, naive T cells, regulatory T cells; CD4 for CD4 T_helper cells, memory T cells, naive T cells, regulatory T cells; CD8A for cytotoxic T cells and naive cytotoxic T cells. We evaluated the performance of predicting a cell type (e.g. B cell) based on the imputed expression of a marker gene (e.g. CD19). For each cell type and marker gene pair, we calculated the area under the receiver operating characteristic (ROC) curve (AUROC) using the `performance` function in the `ROCR` R package where the expression of the marker gene is the predictor and the true cell type is the label. Specifically, the cells were first sorted in a descending order according to the imputed values of the marker gene (e.g. CD19). Consider the cell type *A*. Assume there were *K* cells in total and *b* of them were in cell type *A*. Assume in the top *N* cells, *a* of them were in cell type *A*; in the remaining *K − N* cells, *c* of them were not in cell type *A*. Sensitivity was calculated as *a/b* and specificity was calculated as *c/*(*K − b*). The ROC curve was obtained by plotting sensitivity against 1-specificity for different *N*s (*N* = 1, 2, …, *K*).

### 5.6 Evaluation of imputation on unsupervised clustering

We used two sets of datasets for this analysis. The first set is CellBench^18^ data, which consists of 7 datasets. Three of these datasets contained three cell lines (datasets: *sc_10x*, *sc_dropseq*, *sc_celseq2*) and four datasets contained five cell lines (datasets: *sc_10x_5cl*, *sc_celseq2_5cl_p1*, *sc_celseq2_5cl_p2*, *sc_celseq2_5cl_p3*). The second set of data contains sorted PBMCs3 (*N*=59620) with 10 cell types.

Clustering was performed using both *k*-means^45^ and Louvain clustering^46^ where the number of clusters was set to be the number of known cell types known in each dataset. In *k*-means we directly set *k* to be the number of known cell types, while in Louvain clustering this is achieved by increasing the number of nearest neighbors iteratively until the desired number of clusters is obtained. Louvain clustering was performed by performing feature selection using highly variable genes (HVGs), applying PCA by `prcomp()`, and then building a shared *k*-nearest-neighbors (*k*NN) graph^46^ based on the Euclidean distances of the top 10 PCs. An edge was drawn between all pairs of cells sharing at least one neighbor, weighted by the characteristics of the shared nearest neighbors. For this last step, we used the `buildSNNGraph` function in the `scran`^38^ R/Bioconductor package with the top 10 PCs as input, *d* = *NA* and other parameters as default. Clusters were identified by a multi-level modularity optimization algorithm for finding community structure^59^ using `cluster_louvain` in `igraph` R package. *K*-means clustering was performed using the top 10 PCs.

To evaluate the performance of each method, we used four metrics:

1. Entropy of accuracy (ECA or *H_accuracy_*) where 0 ≤ *H_accuracy_* ≤ log *M*. *H_accuracy_* measures the diversity of the ground-truth labels within each predicted cluster group assigned by the clustering algorithm.

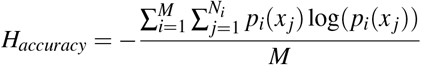

where *M* is the total number of predicted clusters from the clustering algorithm, *N* is the number of ground-truth clusters, and *N_i_* is the number of ground-truth clusters in the *i^th^* predicted cluster. *x_j_* are cells in the *j^th^* ground-truth cluster, and *p_i_*(*x_j_*) are the proportions of cells in the *j^th^* ground-truth cluster relative to the total number of cells in the *i^th^* predicted cluster. A smaller value of *H_accuracy_* is better as it means the cells in a predicted cluster are homogeneous and from the same group^18^. However, *H_accuracy_* can lead to over-clustering, with an extreme case being treating each cell as a cluster (or *H_accuracy_* = 0).
2. Entropy of purity (ECP or *H_purity_*) where 0 ≤ *H_purity_* ≤log *N*. *H_purity_* measures the diversity of the predicted cluster labels within each of the ground-truth groups.

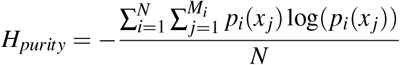

where *N* is the total number of ground-truth clusters, *M_i_* is the number of predicted clusters in the *i^th^* true cluster. *x_j_* are cells in the *j^th^* predicted cluster, and *p_i_*(*x_j_*) are the proportions of cells in the *j^th^* predicted cluster relative to the total number of cells in the *i^th^* ground-truth cluster. A smaller value of *H_purity_* is better as it means the cells in the ground-truth groups are homogeneous with the same predicted cluster labels^18^. However *H_purity_* can lead to under-clustering, with an extreme case being assigning all cells into one predicted cluster so that each of the ground-truth groups has the same predicted cluster label (*H_purity_* = 0).
3. Adjusted Rand index^60^ (ARI). We used the `adjustedRandIndex` function in `mclust` package^61^. The minimum ARI of each dataset was obtained by permuting cells’ ground-truth cell type labels 10^4^ times, recomputing ARI, and averaging the 10^4^ ARIs from random permutations. The maximum has been theoretically proved as 1. We subtract the ARI by the empirical minimum and divided by the distance between the empirical minimum and theoretical maximum.
4. Median Silhouette index *S* = *Median*(*s*(*i*)) where *i* is a cell, *C_i_* is the set of cells in the same cluster as *i*, |*C_i_*| is its cardinality (i.e., number of cells in a cluster),

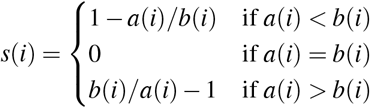

and 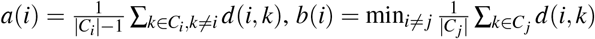. Here *d*(*i, k*) denotes the Euclidean distance between cells *i* and *k*. This is implemented by `silhouette` function in the `cluster` R package^62^. The range of S is [−1, 1].

### 5.7 Identify cell types in *HCA_10x_tissue* dataset

To computationally identify the cell types in the *HCA_10x_tissue* dataset, we downloaded bulk RNA-seq profiles from 13 primary hematopoietic cell types (GEO accession number GSE74246). The data were normalized and scaled using log 2-transformed TPM. We averaged bulk profile replicates from the same cell type. We used the bulk profiles to computationally assign a cell type label for each individual cell. Consider the *j^th^* bulk cell type profile. We calculated the log-fold change (LFC) between the *j^th^* bulk cell type profile and each of the other bulk profile and identified the top 100 genes with the largest LFC for each comparison, which were used to define anchor gene sets. Repeating this for all bulk cell types resulted in 13 × 12 = 156 anchor gene sets. Next, for each bulk cell type or each single cell, we calculated mean expression of anchor genes in each of these gene sets, yielding a vector of length 156 which we refer to as “anchor profile” of a cell type or a cell. We calculated the Spearman correlation between each single cell anchor profile and each bulk cell type anchor profile. A cell was computationally assigned to the *j^th^* cell type if the bulk profile of this cell type had the largest correlation with the single cell and the correlation coefficient was greater than 0.6. If no cell type label was assigned to a cell in this way, the cell’s cell type was labeled as unknown. Next, *k*-means clustering was performed on the cells. For each cluster, if at least 70% of cells were inferred as cell type *A*, then all cells in the cluster were relabeled as cell type *A*. If a cluster cannot be labeled in this way, then all cells in the cluster were labeled as unknown cell type. The largest cluster obtained in this procedure contained *N*=197 cells inferred as monocytes. They were used for the null differential analysis described before.

### 5.8 Evaluation on pseudotime inference

#### 5.8.1 Constructing trajectories

We used the (1) CellBench^18^ RNA mixture (*RNAmix_celseq2*, *RNAmix_sortseq*) and cell mixture (*cellmix1*, *cellmix2*, *cellmix3*, *cellmix4*) datasets from three cell lines (H2228, H1975, HCC827) generated by CEL-seq2 and SORT-seq protocols^18^ and (2) bone marrow cells from the HCA_10x_tissue^11^. Monocle 2^50^ and TSCAN^49^ were used to construct trajectories on the imputed expression values. Monocle 2 uses a DDR-Tree (Discriminative DRTree)^63^ for dimensionality reduction and tree construction. For TSCAN, we calculated the top principal components (PCs) with the `prcomp` function in the `stats` R package. We then run *k*-means clustering (using the `kmeans` function in the `stats` R package) on the top PCs (obtained by TSCAN using elbow method) to obtain predicted cluster labels for each cell. Then, we used the predicted cluster labels and top PCs as input to TSCAN. The number of clusters was chosen to be the smallest one that allowed two branches in the spanning tree to be consistent with the underlying true tree structure.

To identify the root cells or root state when inferring trajectories with TSCAN and Monocle 2 using CellBench data, the cluster with the most H2228 cells was selected. For *HCA_10x_tissue data*, each cell was assigned a differentiation level, as follows. HSC: level 1; MPP: level 2; LMPP and CMP: level 3; CLP, GMP, and MEP: level 4; B cell, CD4 T cell, CD8 T cell, NK cell, Monocyte, and Erythroid: level 5. Then, the cluster of cells with the smallest averaged differentiation level was set to be root state for the inferred trajectories.

#### 5.8.2 Assessment

For the CellBench^18^ data, we used the same performance metrics as in Tian et al.’s work^18^: (1) the Pearson correlation between the inferred trajectory and the rank order of the cells for which we know the true ordering of the cells, and (2) the proportion of cells on an inferred trajectory for which the inferred branch and the true branch were the same.

For the bone marrow cells from the *HCA_10x_tissue data*, we compared the estimated pseudotime to the level of differentiation for each pair of cells. For instance, for a pair of cells A and B, if cell A is a HSC (differentiation level 1), cell B is a MPP (differentiation level 2), and the inferred pseudotime for cell A and B is *t_A_* and *t_B_* where *t_A_* < *t_B_*, then we call it “correctly ordered”. The *percentage* of the correctly ordered cells averaged across all possible cell pairs served as the performance metric.

### 5.9 Performance of imputation algorithms on time, memory and scalability

We created four datasets for this analysis. Using the *10x_293t_jurkat* dataset, we created two smaller datasets by randomly sampling *N*=1000 and *N*=5000 cells (*1k_cell*, *5k_cell*, respectively). Using the *HCA_10x_tissue* dataset, we created two larger datasets by randomly sampling *N*=50,000 and *N*=100,000 cells (*50k_cell*, *100k_cell*, respectively). By running the imputation methods on all datasets, we assessed the computational time (in minutes), memory usage (in maximum resident set size of all tasks in job (MaxRSS or maximum resident set size of all tasks in a job) in gigabytes returned from the Slurm command `sacct`), and scalability with respect to cell number. For each method, a linear model was fit using the `lm` function from the `stats` R package where the computation time was the response and the number of cells on the log_10_-scale was the predictor (Figure S9). The coefficient of the cell number represents the scalability of the method. For methods that failed to produce results, the running time was set to be the maximum time (72 hours) plus 1 minute, and the memory was set to be the maximum memory. The time and memory were linearly scaled to [0, 1]. The average scaled time and memory across all datasets were used as the final score as shown in Fig.6.

### 5.10 Overall performance score

All the assessment measures were mapped to [0, 1] by subtracting the theoretical minimum and then dividing by the difference of the theoretical maximum and minimum. Empirical extrema were used when theoretical ones did not exist. Ten thousand permutations was applied to obtain the empirical minimum of ARI for each dataset. *H_accuracy_* and *H_purity_* were further subtracted by 1 for the convenience of applying “the higher the score, the better the performance” criterion. Efficiency measures (time, memory, scalability) were scaled with the extrema of all methods’ performance.

Evaluation of imputed values through the similarity to bulk samples, differential analysis, clustering and pseudotime inference were considered as four main assessment aspects. In each aspect, the scores across datasets were first averaged, and next the mean from different analysis tools (MAST and Wilcoxon; *k*-means and Louvain clustering; Monocle 2 and TSCAN) were averaged, then the mean of multiple assessment statistics (e.g. overlap and correlation in assessing pseudotime inference) was used as the evaluation score. The mean scores of all four assessment aspects was finally used to rank all methods.

### 5.11 Data and code availability

The data used in this analysis are described in Section 5.1 and Table S1 with all links or GEO accession numbers. The imputation methods are described in Table S2. All code to reproduce the presented analyses are available at https://github.com/Winnie09/imputationBenchmark. The R package `ggplot2`^64^ for data visualization was used.

## Acknowledgement

This work is supported by the National Institutes of Health grants R01HG010889 and R01HG009518 to HJ and R00HG009007 to SCH. SCH is also supported by the Chan Zuckerberg Initiative DAF grants 183201 and 356-01. WH and SCH are supported by Alex’s Lemonade Stand Foundation. We would like to thank the Maryland Advanced Research Computing Center (MARCC) for providing computing resources.

## Supplementary

### Supplementary Tables

**Table S1. Summary of all datasets used in each evaluation of this benchmark**. The table includes the names, protocols, source (GEO accession numbers or links to download) and cell details of each dataset.

**Table S2. Summary of all scRNA-seq imputation methods used in each evaluation of this benchmark**. The table includes the name of the method, input, output, pre-processing steps for each method that we applied, the programming language, assumptions about the me thod, the download date, software version number, and link to software package.

**Table S3. Values of all three efficient measures in time, memory and scalability using all four datasets.** The table include the computation time and memory of four datasets with 10^3^, 5 × 10^3^, 5 × 10^4^, 10^5^ cells for all imputation methods. Scalability is the coefficient of the cell number of each dataset in the linear model where the number of cells on the log_10_-scale is fitted against the computation time.

### Supplementary Figures

**Figure S1.**
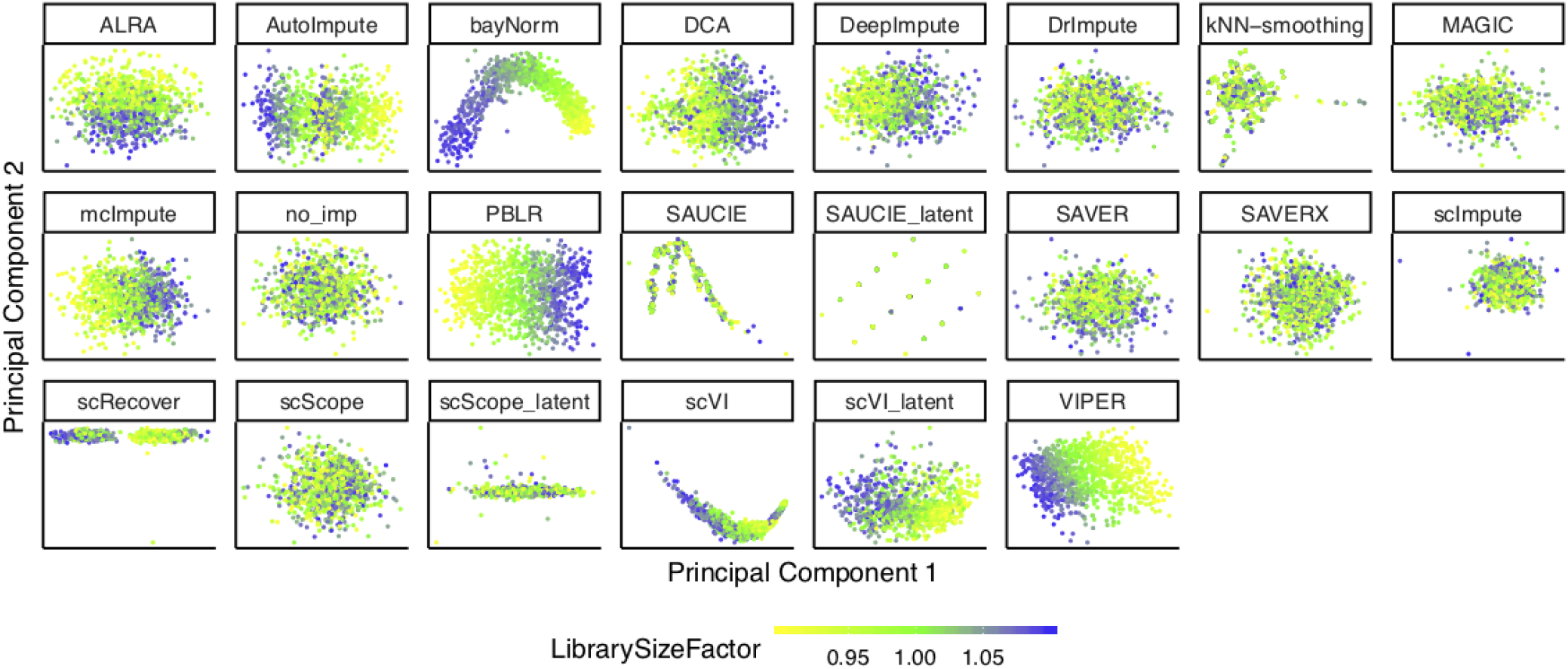
Dimension reduction results after applying Principal Components Analysis (PCA) from either no imputation (*no_imp*) or the 18 imputation methods using the *null simulations* data (Section 5.3), except the difference between this figure and Figure 1A is this figure includes the latent spaces directly found by scScope (scScope_latent), scVI (scVI_latent) and SAUCIE (SAUCIE_latent) (not Principal Components – PCs). All other methods are showing observations along the first two PCs. The color represents the simulated library size (defined as the total sum of counts across all relevant features) for each cell.

**Figure S2.**
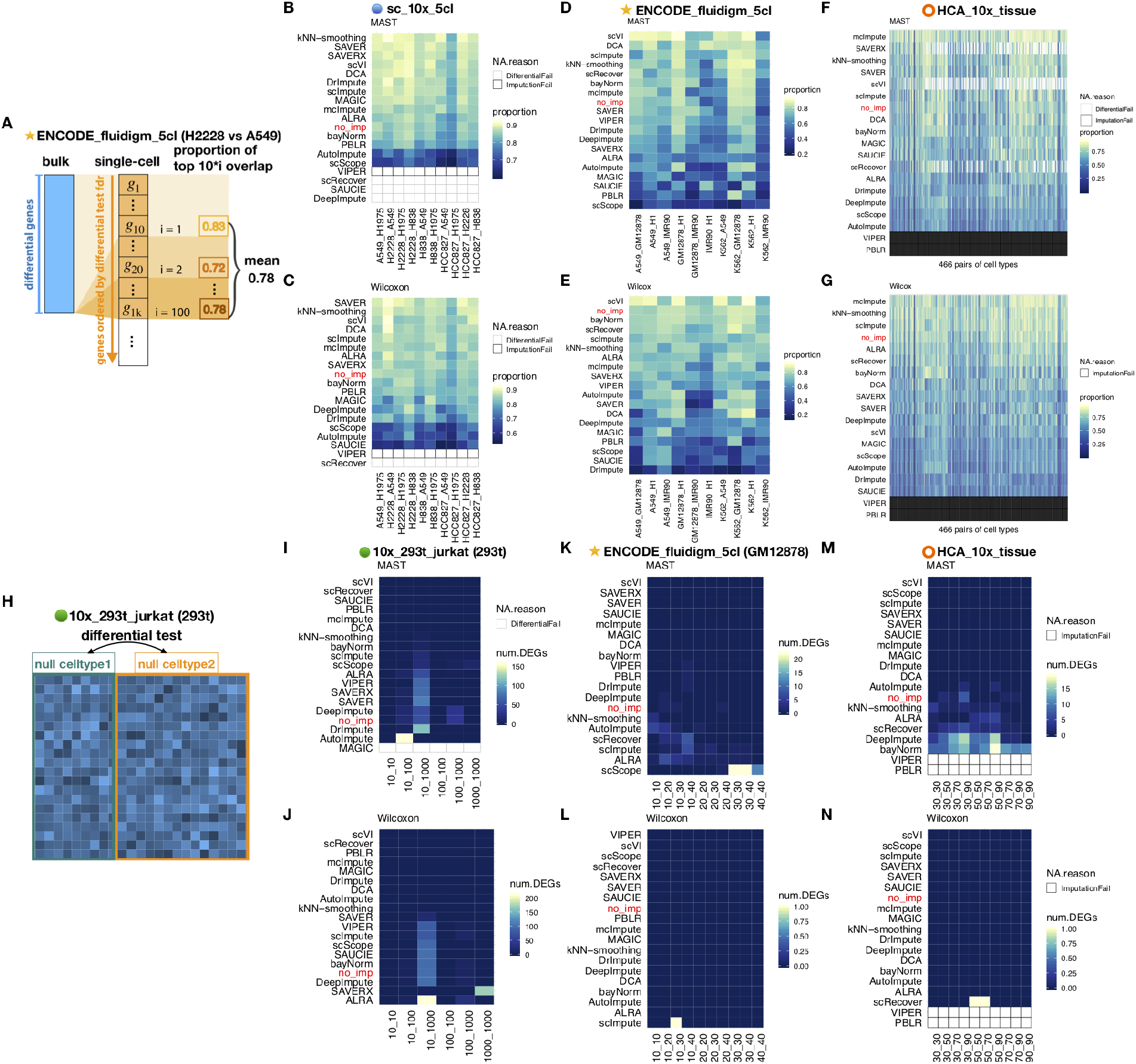
Impact of imputation methods on differential expression analysis. **(A)** Schematic of evaluating differentially expressed genes (DEGs) using the overlap between bulk RNA-seq and scRNA-seq – also shown in Figure 3A. Using the pairs of cell lines in the *sc_10x_5cl* dataset, *ENCODE_fluidigm_5cl* dataset, and pairs of cell types in the bone marrow tissue from the *HCA_10x_tissue* dataset, we show heatmaps of proportion of overlap between bulk and single-cell DEGs identified using **(B, D, F)** MAST^43^ and **(C, E, G)** Wilcoxon-rank-sum test^44^ (abbreviated as Wilcoxon) for differential expression, respectively. **(H)** Schematic of a null differential expression analysis by randomly partitioning cells from the same cell type into two groups – also shown in Figure 3E. Using the 293T cells from the *10x_293t_jurkat* dataset, the GM12878 cells from the *ENCODE_fluidigm_5cl* dataset, and bone marrow cells from the *HCA_10x_tissue* dataset, the number of false positive DEGs identified using **(I, K, M)** MAST and **(J, L, N)** Wilcoxon, respectively. The x-axis in Figures (I-N) describe the number of cells in each group (e.g. 10 sampled cells in group 1 and 10 sampled cells in group 2) when applying a method to identify differentially expressed genes. White areas with black outline indicate that the imputation methods did not return output after 72 hours and areas with grey outline indicate that either MAST or Wilcoxon failed to return results.

**Figure S3.**
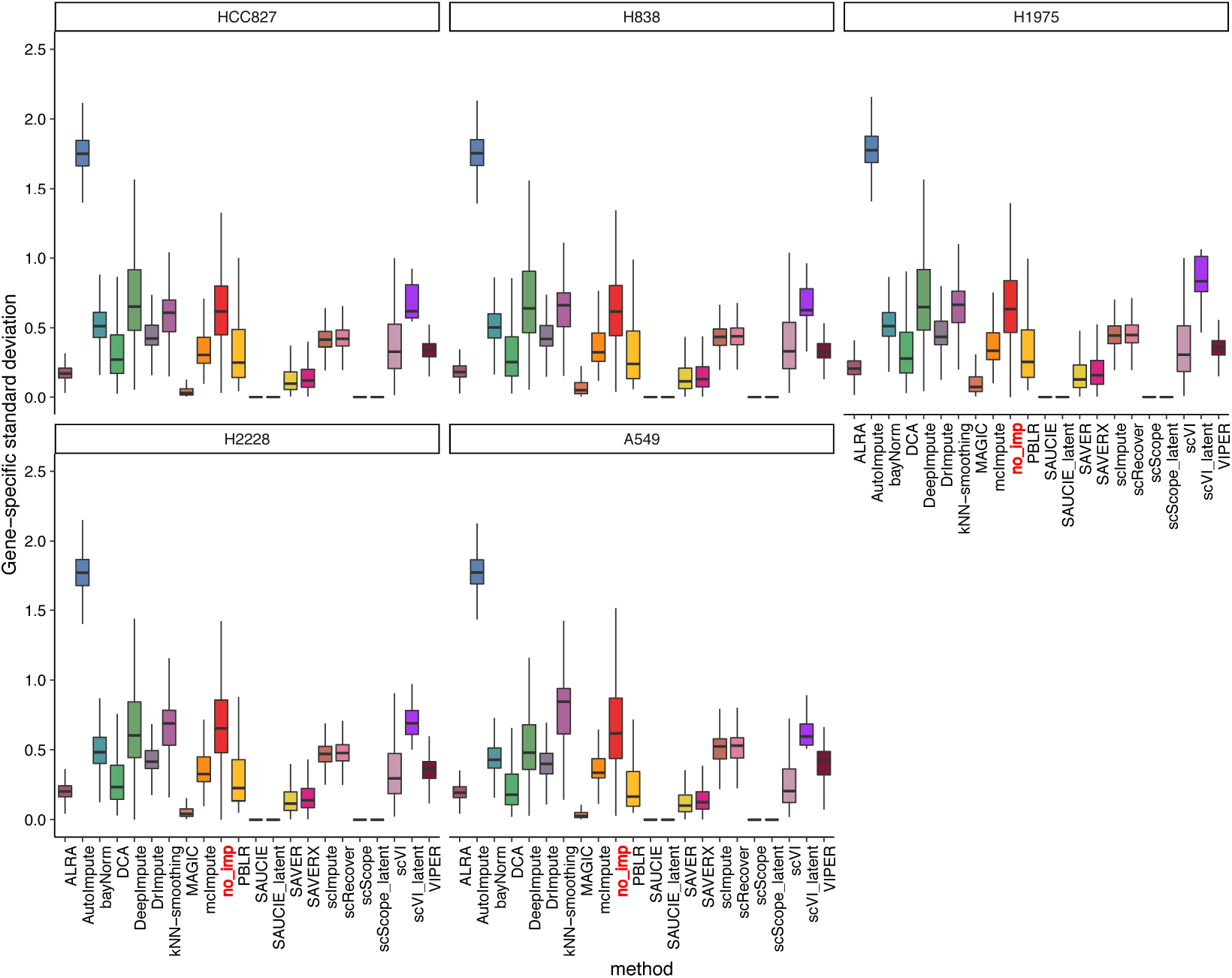
Estimates of gene-specific standard deviation of imputed values for five cell lines. For each cell line in the *sc_10x_5cl* dataset, we calculated the gene-specific standard deviation across cells with no imputation (*no_imp*) and with imputation.

**Figure S4.**
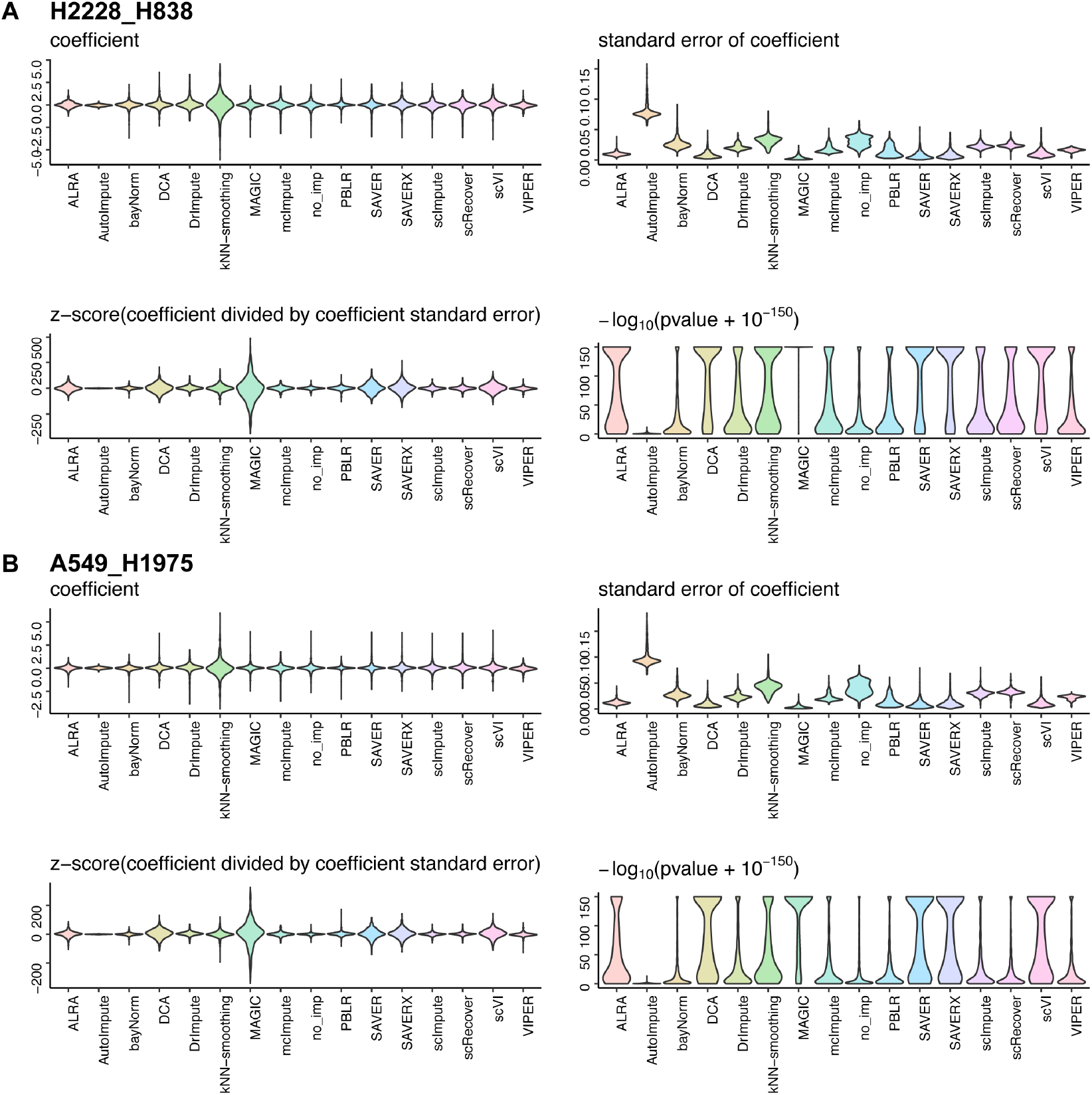
Summary distributions of output from MAST using the imputed values from pairs of cell lines. The pairs of cell lines from the *sc_10x_5cl* dataset compared below are **(A)** H2228 (*N*=758) vs H838 (*N*=876) – more balanced group sizes and **(B)** A549 (*N*=1256) vs H1975 (*N*=440) – less balanced group sizes. For each pair of cell lines, we applied MAST and extracted and plot the following information: the distribution of estimated log-fold changes (‘coefficients’) (top left), the distribution of standard errors for the coefficients (top right), the distribution of test statistics (z-score output from MAST, bottom left), and distribution of log-transformed *p*-values (bottom right).

**Figure S5.**
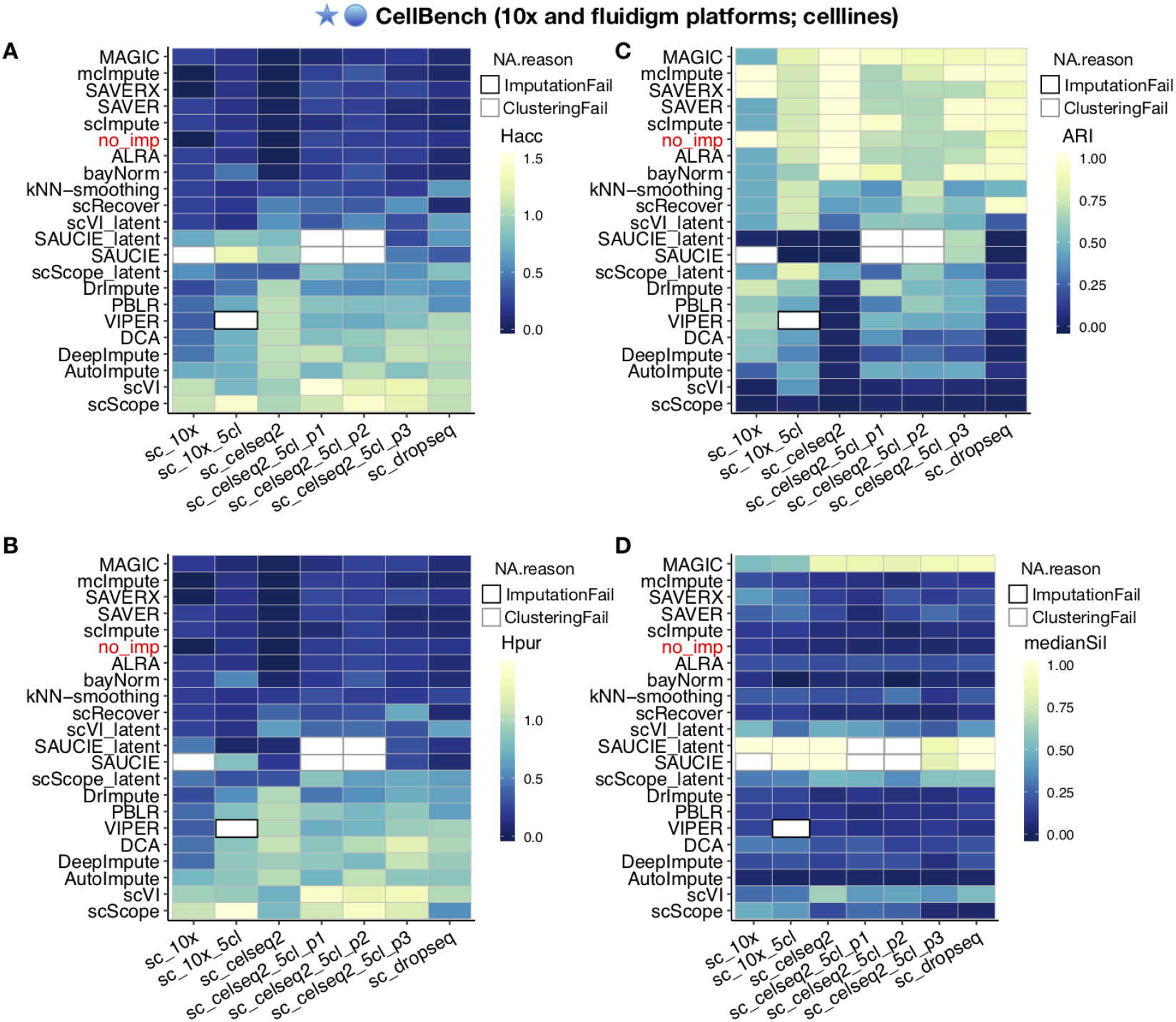
Impact of imputation methods on *k*-means clustering analysis using seven datasets from CellBench single cells. Heatmaps of the individual performance metrics **(A)** entropy of cluster accuracy (*H_acc_*), **(B)** entropy of cluster purity (*H_pur_*), **(C)** adjusted Rand index (ARI), and **(D)** the median of Silhouette and median Silhouette index of each imputation method for each of the seven datasets in CellBench^18^. The white boxes with black lines represent cases in which no output was returned from the imputation method after 72 hours. The white boxes with gray lines represent cases in which the clustering algorithm failed to cluster the cells using the principal components, for instance “more cluster centers than distinct data points” because many cells with imputed profiles are identical.

**Figure S6.**
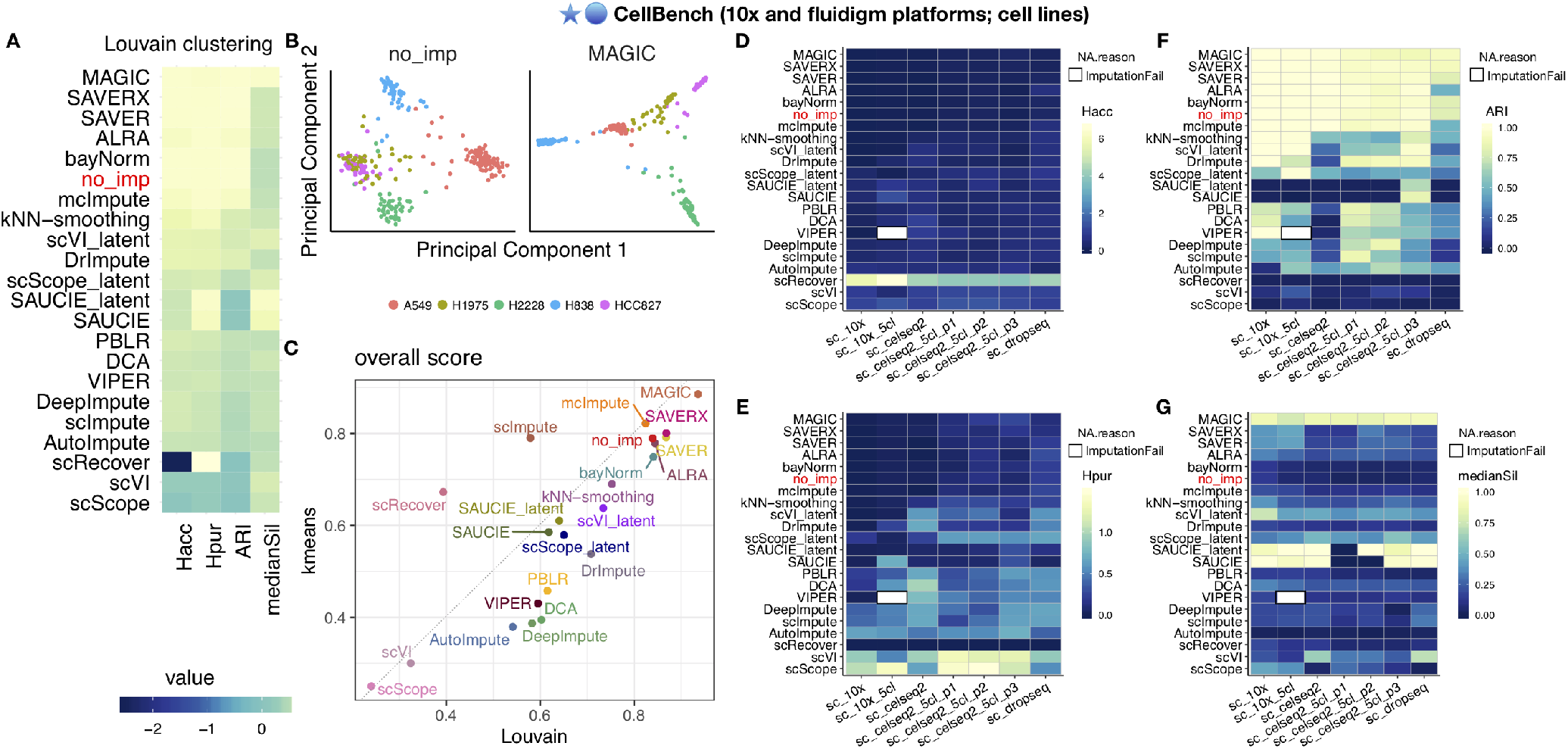
Impact of imputation methods on Louvain clustering analysis using seven datasets from CellBench^18^. **(A)** Heatmap of four performance metrics – entropy of cluster accuracy (*H_acc_*), entropy of cluster purity (*H_pur_*), adjusted Rand index (ARI), and median Silhouette index – averaged across seven datasets. To compare imputation methods across metrics, the metrics were re-scaled to be between 0 and 1 and the order of *H_acc_* and *H_pur_* were flipped to where a higher standardized score translates to better performance. Imputation methods (rows) are ranked by the average across all four metrics. **(B)** Dimension reduction results after applying PCA to the *sc_celseq2_5cl_p1* data with no imputation (left) and with imputation using MAGIC (right). The colors are the true group labels. **(C)** Overall score (or average of the four performance metrics) for Louvain clustering (x-axis) and *k*-means clustering (y-axis). **(D-G)** Heatmaps of the individual performance metrics (D) *H_acc_*, (E) *H_pur_*, (F) *ARI* and (G) the median of Silhouette of each imputation method for each CellBench dataset. The white boxes with black lines represent cases in which no output was returned from the imputation method after 72 hours.

**Figure S7.**
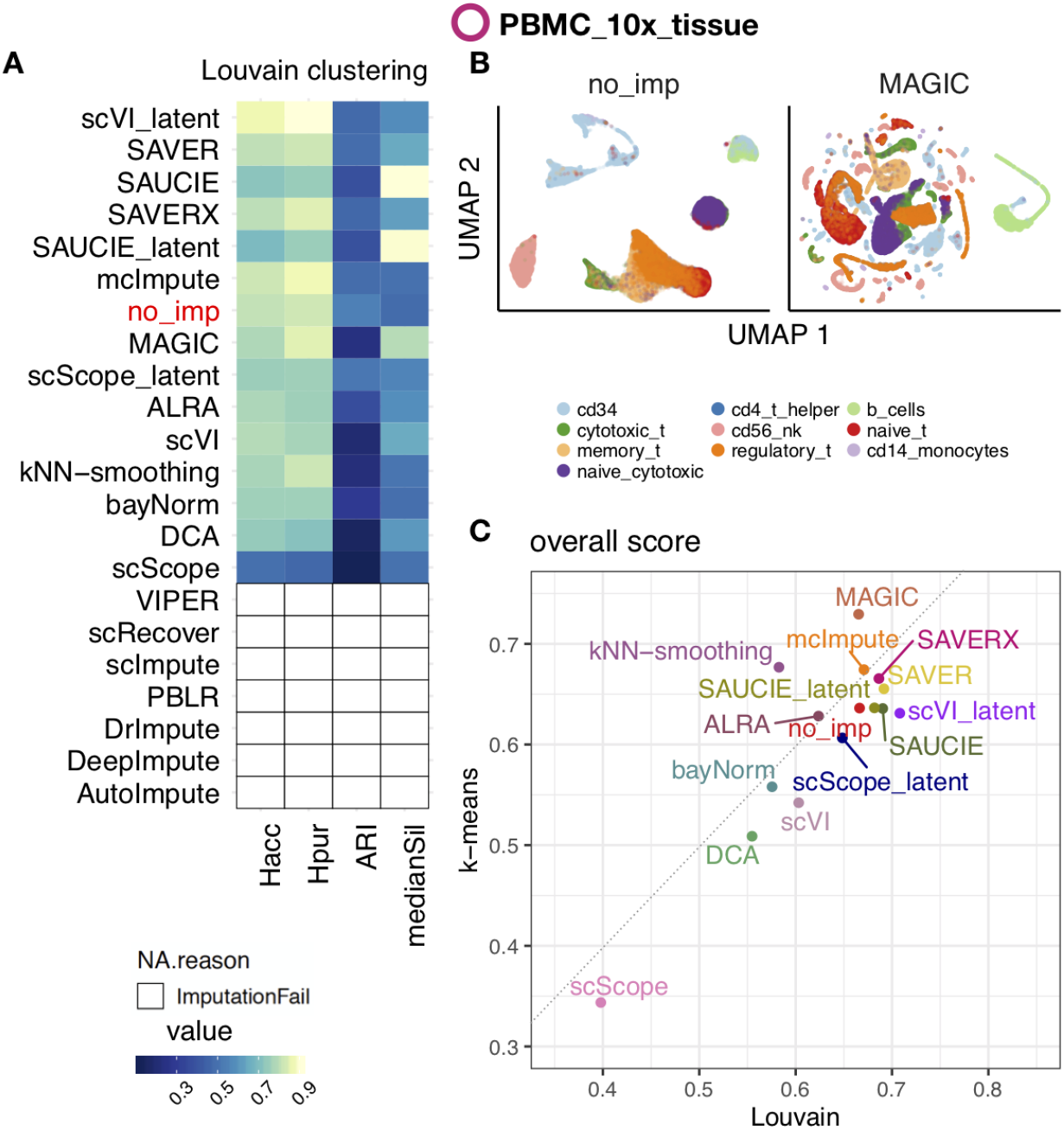
Impact of imputation methods on Louvain clustering analysis using ten sorted peripheral blood mononuclear cell (PBMC) cell types from 10x Genomics. **(A)** Heatmap of four performance metrics – entropy of cluster accuracy (*H_acc_*), entropy of cluster purity (*H_pur_*), adjusted Rand index (ARI), and median Silhouette index – on data from 10x Genomics^3^. To compare imputation methods across metrics, the metrics were re-scaled to be between 0 and 1 and the order of *H_acc_* and *H_pur_* were flipped to where a higher standardized score translates to better performance. Imputation methods (rows) are ranked by the average across all four metrics. **(B)** Dimension reduction results using UMAP components^48^ with no imputation (left) and with imputation using MAGIC (right). The colors are the true group labels. **(C)** Overall score (or average of the four performance metrics) for Louvain clustering (x-axis) and *k*-means clustering (y-axis). White areas with black outline in (D) indicate that the imputation methods did not return output after 72 hours.

**Figure S8.**
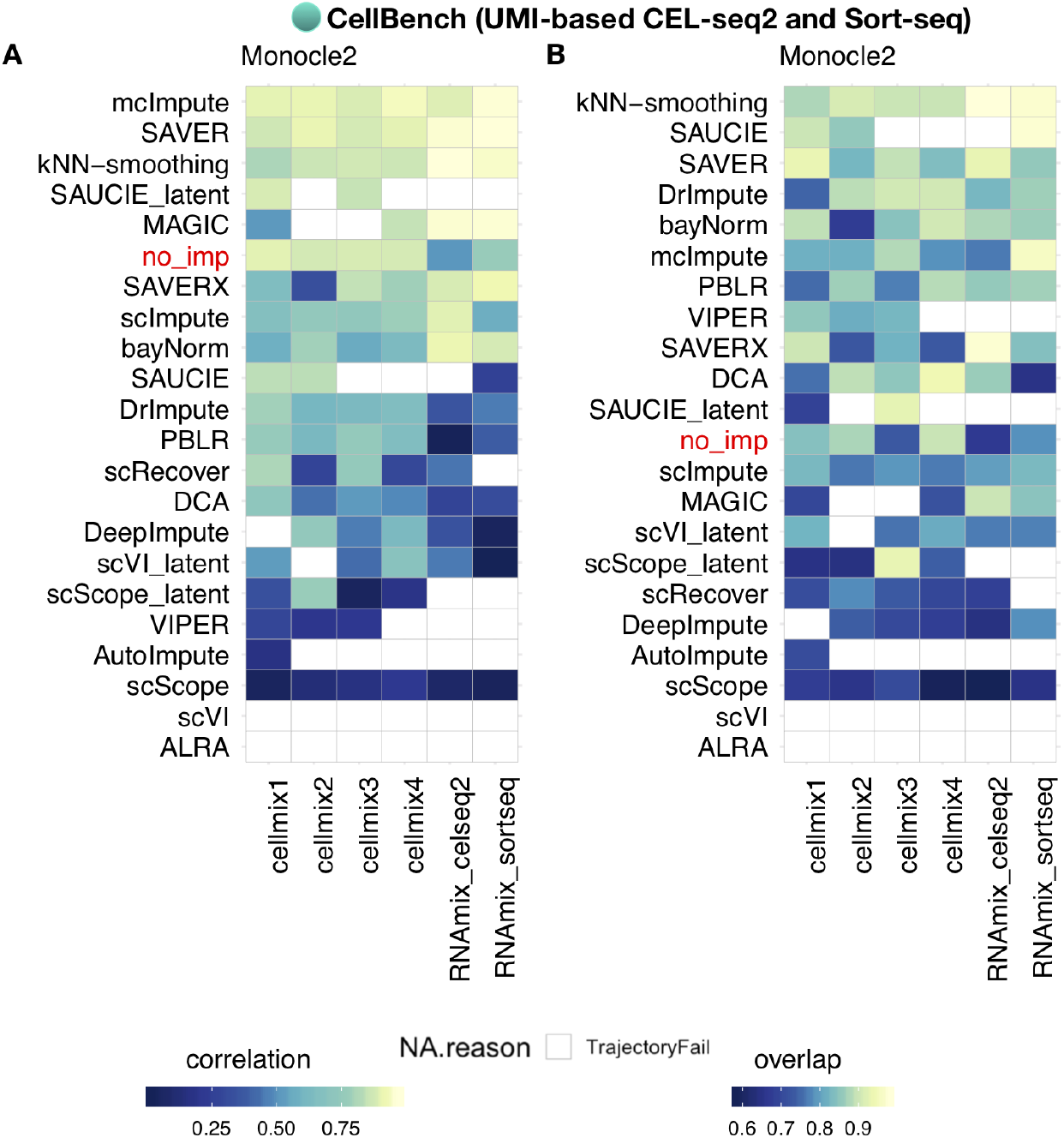
Impact of imputation methods on inferred trajectories for pseudotime analysis using Monocle 2 with the RNA mixture and cell mixture datasets from CellBench^18^. **(A)** Heatmap showing the Pearson correlation coefficients (PCC), denoted as *correlation*, between the inferred trajectory and the rank order of the cells where we know the true trajectory (or ordering) of the cells. **(B)** Heatmap of the proportion of cells on the inferred trajectories that correctly *overlap* with the cells on the branch where we know the true trajectory of the cells.

**Figure S9.**
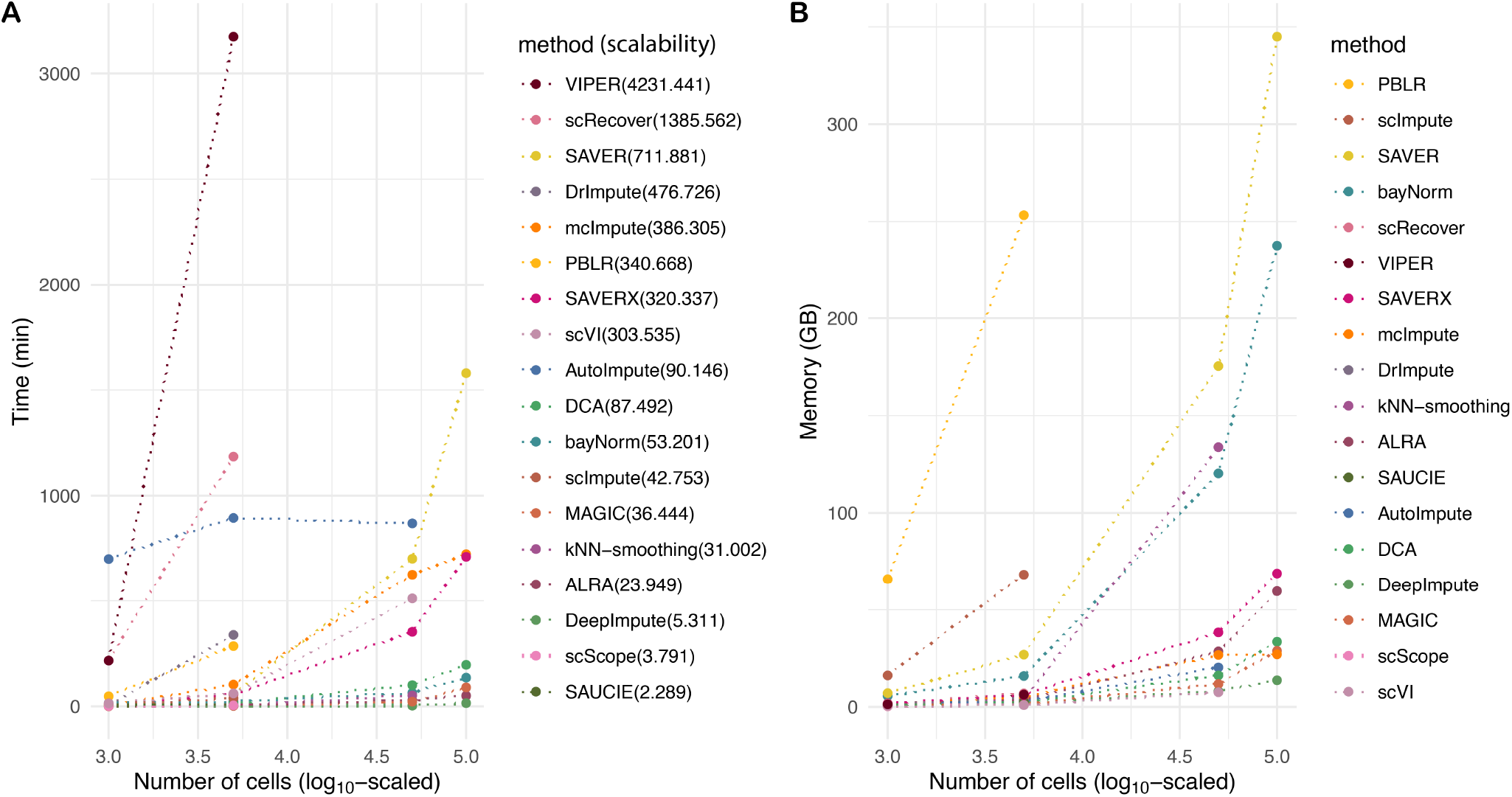
Values of time, scalability and memory of each imputation method on each dataset before scaling. **(A)** Computation time (in minutes) for each method to finish imputing four datasets of 10^3^, 5 × 10^3^, 5 × 10^4^ and 10^5^ cells, respectively. For some imputation methods, no results are shown because no imputed values were returned within 72 hours. Scalability (marked in the parentheses after each methods’s name) is defined by fitting a linear model with the number of cells on the log_10_-scaled on the x-axis and the computation time on the y-axis and using the coefficient as the metric for scalability. **(B)** Memory usage (in maximum resident set size of all tasks in job (MaxRSS) in gigabyte, i.e. GB, returned from the Slurm command `sacct`) for each method to finish imputating the four datasets mentioned above.

